# Systematic analysis of cellular crosstalk reveals a role for SEMA6D-TREM2 regulating microglial function in Alzheimer’s disease

**DOI:** 10.1101/2022.11.11.516215

**Authors:** Ricardo D'Oliveira Albanus, Gina M Finan, Logan Brase, Shuo Chen, Qi Guo, Abhirami Kannan, Mariana Acquarone, Shih-Feng You, Brenna C Novotny, Patricia M Ribeiro Pereira, John C Morris, David M Holtzman, Eric McDade, Martin Farlow, Jasmeer P Chhatwal, Dominantly Inherited Alzheimer Network (DIAN), Emily E Mace, Bruno A Benitez, Laura Piccio, Greg T Sutherland, Qin Ma, Hongjun Fu, Celeste M Karch, Oscar Harari, Tae-Wan Kim

**Author notes:** **Contributed equally** (Ricardo D'Oliveira Albanus, Gina M Finan).

## Abstract

Cellular crosstalk, mediated by membrane receptors and their ligands, is crucial for brain homeostasis and can contribute to neurodegenerative diseases such as Alzheimer’s disease (AD). To discover crosstalk dysregulations in AD, we reconstructed crosstalk networks from single-nucleus transcriptional profiles from 67 clinically and neuropathologically well-characterized controls and AD brain donors. We predicted a significant role for TREM2 and additional AD risk genes mediating neuron-microglia crosstalk in AD. The gene sub-network mediating SEMA6D-TREM2 crosstalk is activated near Aβ plaques and *SEMA6D*-expressing cells and is disrupted in late AD stages. Using CRISPR-modified human induced pluripotent stem cell-derived microglia, we demonstrated that SEMA6D induces microglial activation in a *TREM2*-dependent manner. In summary, we demonstrate that characterizing cellular crosstalk networks can yield novel insights into AD biology.

**One Sentence Summary:** We investigate cell-to-cell communication in Alzheimer’s disease to characterize disease biology and suggest new avenues for therapeutic intervention.

## INTRODUCTION

Cross-cellular signaling (cellular crosstalk) is integral to normal brain physiology. By establishing cellular networks mediated by membrane receptors and their corresponding ligands, cells can gather information from their immediate environment and respond accordingly. Indeed, cellular crosstalk is crucial to brain homeostasis and processes of neurodevelopment, such as synaptic pruning and axon guidance *(1, 2)*. However, increasing experimental and genetic evidence implicates aberrant cellular crosstalk as a contributing factor to neurodegenerative diseases, including Alzheimer’s disease (AD) *(3–6)*. Furthermore, from a translational perspective, cellular crosstalk is an attractive molecular target for drug development, as membrane receptors are relatively amenable to therapeutic targeting *(7–9)*. Therefore, systematic characterization of brain cellular crosstalk interactions can help identify molecular mechanisms involved in neurodegeneration and inform novel therapies.

Genome-wide association studies (GWAS) have successfully identified genetic risk loci for AD and nominated genes likely mediating these genetic signals *(10–13)*. Further, by leveraging human tissue *(14–17)* and experimental data from human induced pluripotent stem cell (iPSC)- derived cells *(18)*, functional genomics studies have revealed that many AD risk genes are expressed by microglial cells. However, how most of these microglial AD risk genes are regulated in the contexts of normal physiology and AD pathophysiology is still unknown. As the resident immune cells of the brain, microglia are highly attuned to their surrounding environment, including signals from neighboring cells *(19)*. While previous studies have shown causal effects of disrupted crosstalk in neurodegeneration, it remains unclear how cellular crosstalk between microglia and other cell types is involved in mediating AD genetic risk.

Understanding these processes requires techniques that can systematically characterize the crosstalk networks in the brain, reconstruct the likely signaling pathways downstream of these interactions, and integrate these data with genetic findings.

In this study, we used single-nucleus transcriptomic profiles (snRNA-seq) from clinically, neuropathologically, and genetically well-characterized human brains to systematically reconstruct the cellular crosstalk networks across seven major brain cell types: microglia, astrocytes, oligodendrocytes, oligodendrocyte precursors (OPCs), excitatory and inhibitory neurons, and endothelial cells. We found that direct involvement of known AD risk genes was more frequent in neuron-microglia crosstalk interactions than in those between other cell types. In addition, we identified a sub-network of microglial genes centered around *TREM2* that we predicted mediates neuron-microglia crosstalk. We predicted that this sub-network is modulated by the crosstalk interaction between neuronal semaphorin 6D (SEMA6D) and microglial TREM2. We found evidence that this sub-network is disrupted in late-stage AD, and, using spatial transcriptomics, we observed that the *TREM2* sub-network is activated in proximity to Aβ plaques and *SEMA6D*-expressing cells. Finally, we validated our predictions *in vitro* using wild-type (WT) and *TREM2* knockout (KO) human iPSC-derived microglia (iMGL). We showed that SEMA6D promotes microglia functions, including phagocytosis and cytokine release, in a *TREM2*-dependent manner. Our findings demonstrate that systematic characterization of cellular crosstalk networks in human brains is a viable strategy to elucidate aberrant regulatory biology in AD and other neurodegenerative diseases, which could ultimately inform the development of novel AD therapies.

## RESULTS

### A complex landscape of crosstalk dysregulation in AD

To systematically characterize cellular crosstalk interactions in controls and AD, we analyzed snRNA-seq profiles of superior parietal cortex tissue samples from brain donors of the Knight Alzheimer Disease Research Center (Knight ADRC) and the Dominantly Inherited Alzheimer Network (DIAN), previously published by our group *(14)*. This dataset encompasses different AD subtypes, including sporadic AD and autosomal dominant AD, with donors distributed in a broad spectrum of neuropathological states and genetic backgrounds, including carriers of *TREM2* risk variants **(table S1)**. In total, we analyzed ∼300K nuclei representing seven major brain cell types (microglia, astrocytes, oligodendrocytes, OPCs, excitatory and inhibitory neurons, and endothelial cells) from 67 donors (**Fig. 1A**). We identified patterns of ligand-receptor gene expression across cell type pairs using CellPhoneDB *(20)*, which has been successfully used to predict patterns of brain cellular crosstalk *(5)*.

**Fig. 1:**
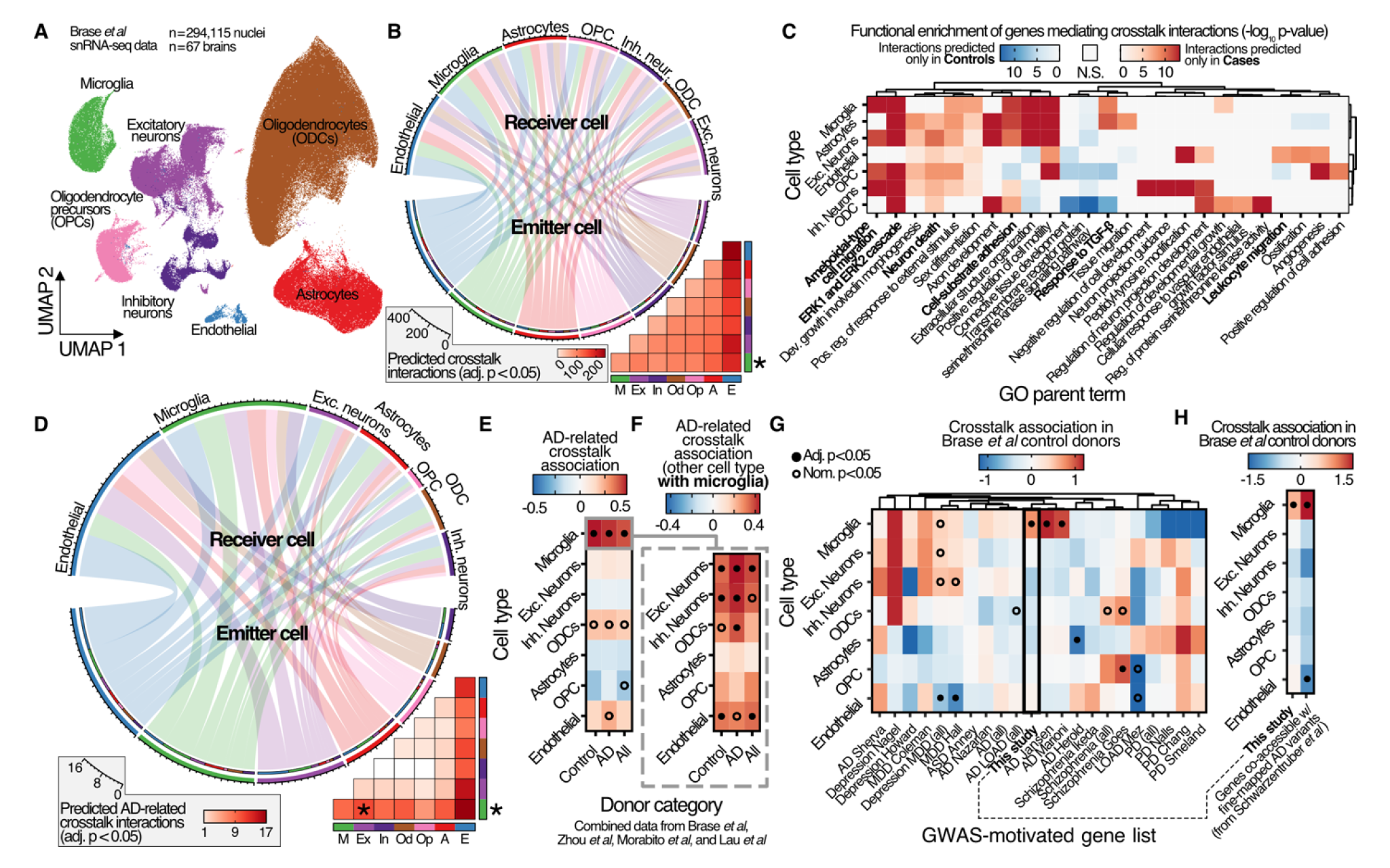
Overview of predicted cellular crosstalk interactions in human brains. **A)** Uniform Manifold Approximation and Projection (UMAP) representation of the snRNA-seq dataset identifying the seven major brain cell types investigated in this study. **B)** Number of unique significant CellPhoneDB interactions detected involving each cell type as either the ligand or receptor across donor categories. Asterisks indicate cell types with significantly different number of predicted crosstalk interactions between cases and controls (from fig. S2). **C)** Gene ontology enrichments of genes mediating crosstalk interactions only detected in cases (red colors) and controls (blue colors). **D)** Cellular crosstalk interactions involving one AD gene as either ligand or receptor across all cellular state pairs. Asterisks indicate cell types (outermost asterisk) and cell type pairs (heatmap asterisk) significantly enriched for AD-related interactions. **E)** Association of AD crosstalk interactions for each cell type using combined data from multiple snRNA-seq datasets. **F)** Association of AD crosstalk interactions between microglia and other cell types. **G)** Crosstalk associations for each cell type (control donors from this study) in genes nominated by GWAS from multiple neuropsychiatric traits. **H)** Similar to (G) but using only AD genes supported by snATAC-seq co-accessibility (Methods).

We predicted crosstalk interactions separately for disease status and genetic group in our CellPhoneDB analyses. In total, we identified between 961 and 1,600 (median = 1,521) significant (Bonferroni-corrected *p* < 0.05) crosstalk interactions between cell type pairs across all donor categories **(Fig. 1B**; **fig. S1A**; **table S2**). We compared the crosstalk patterns across cases and controls to identify global changes associated with disease status. We predicted significantly more crosstalk interactions involving microglia in AD donors compared to controls (odds ratio; OR = 1.12, *p* = 0.019, Fisher’s exact test; **Fig. 1B**; **fig. S2**). Because there is limited evidence on whether microglia numbers change during AD *(21)*, and we did not observe significant alterations in the count of microglial nuclei associated with disease state, we speculate that the increased number of predicted crosstalk interactions involving microglia in AD donors indicates transcriptional changes consistent with microglial activation. This suggests that changes in microglial function and state, rather than changes in cell abundance, result in changes in crosstalk patterns.

We next performed a functional enrichment of the genes involved in the predicted crosstalk interactions specific to each subset of donors to determine which biological pathways are disrupted in AD. Consistent with widespread perturbations of normal brain physiology in AD, we observed changes in the cellular crosstalk patterns when comparing AD donors to controls. The genes involved in crosstalk interactions predicted only in AD donors were significantly enriched for pathways associated with immune activation and migration (*e.g.,* response to transforming growth factor β, ameboid cell/leukocyte migration) and neuronal stress (*e.g.,* neuron death, ERK1/2 cascade; **fig. S3**). Aiming to understand how each cell type likely contributes to the dysregulated pathways in AD, we performed a functional enrichment analysis separately for the crosstalk interactions detected in each cell type in cases versus controls. Importantly, we observed that crosstalk interactions involving other cell types besides microglia and neurons were enriched for the very same immune activation and impaired neuronal homeostasis pathways identified in cases versus controls, further supporting that cellular crosstalk contributes to these core features of neurodegeneration (**Fig. 1C**). Together, these results indicate that AD leads to widespread dysregulation of homeostatic cross-cellular signaling pathways between microglia and neurons with other cells.

### Neuron-microglia crosstalk interactions are enriched to involve known AD risk genes as ligands or receptors

Our initial analysis yielded a vast array of data, predicting thousands of crosstalk interactions across all cell types (**Fig. 1B**). The enormity of this data presented a challenge in discerning the precise role of cellular crosstalk in AD. To render this task more tractable and align it closely with understanding AD biology, we subsequently concentrated our analysis on interactions involving genes empirically linked to AD through genetic and functional studies. We identified 90 possible crosstalk interactions directly involving an AD gene as the ligand or receptor (**table S3**). Of these, 34 were significant by CellPhoneDB analyses in at least one cell type pair (**Fig. 1D**). We calculated for each cell type the association with crosstalk interactions involving AD genes using a logistic regression approach (Methods). Microglia had the highest association for crosstalk interactions involving AD genes across all cell types regardless of which donor subset we analyzed (association range = 0.25 to 0.68, *p* range = 0.015 to 3.53e-3; **fig. S4A**). Notably, most AD gene interactions (64.9%) were predicted to involve microglia as the receptor cell (**table S2**). This observation implies that a subset of AD genes in microglia are modulated by disease-associated alterations in other cells. Therefore, it suggests that the prominent role of microglia in AD may be the endpoint of a cascade initiated within other cell types. This highlights the intricate intercellular complexity of disease pathogenesis and underscores the importance of understanding the role of cellular crosstalk in the development and progression of AD.

To further understand these patterns, we explored the cell types most likely to interact with microglia through crosstalk interactions involving AD genes. We calculated the association of crosstalk interactions involving AD genes for microglia interactions with each cell type. We found that excitatory neurons displayed the highest association with microglia for these interactions (association range = 0.60 to 1.12, *p* range = 0.042 to 1.11e-3; **fig. S4B**). To further validate our findings, we leveraged data from three additional case-control snRNA-seq studies to perform a joint analysis (mega-analysis). These datasets were drawn from the prefrontal cortex region and sourced from the South West Dementia Brain Bank (SWDBB), the Rush ADRC, and the University of California Irvine Institute for Memory Impairments and Neurological Disorders (UCI MIND) ADRC (**table S4**) *(16, 22, 17)*. We observed the same consistent, strong association of microglia with AD-related crosstalk interactions in each study individually or in combination (association range: 0.24 to 0.68, median = 0.53, *p* range: 0.022 to 3.49e-04, median = 6.17e-03; **Fig. 1F, fig. S4A**). We also observed the strongest association of crosstalk interactions involving AD genes for microglia with excitatory neurons (association range = 0.34 to 0.44, *p* range = 4.07e-04 to 3.38e-03; **Fig. 1G**), as well as weaker but significant associations with inhibitory neurons, oligodendrocytes, and endothelial cells. Among the individual studies, we observed some variability concerning which type of broad neuronal cell (excitatory or inhibitory) had the highest association for interactions with microglia involving AD genes (**fig. S4B)**. Together, our findings highlight similar overarching patterns across cohorts and brain regions. This convergence of results suggests that a subset of genes previously linked with AD may facilitate cell signaling pathways between neurons and microglia.

### Cellular crosstalk pattern predictions are robust to cell representation and other potential confounding factors

To determine that our previous results were not driven by the cell type composition of the datasets, the higher representation of AD genes in microglia compared to other cell types, or other possible confounding factors, we statistically controlled for different potential sources of bias in our analyses. First, we observed that the global crosstalk patterns remained similar with dataset downsampling, including removing a subset of donors, using a single donor, and using at most 100 barcodes per snRNA-seq cluster (fraction of replication 0.81 to 0.86, median = 0.84; **fig. S1B-E**). These results indicate that the crosstalk interaction patterns identified using CellPhoneDB are highly robust to the number of donors, skews in donor representation, cell type representation, number of nuclei, and sequencing depth.

Next, we tested whether microglia expressed more genes present in the CPDB database and if this could confound our findings. We observed that microglia and endothelial cells indeed expressed more genes listed as putative ligands or receptors in CellPhoneDB (**fig. S5A**). However, despite more genes associated with crosstalk interactions being expressed in microglia, we did not observe an overrepresentation of AD genes participating in microglia crosstalk interactions compared to other cell types (**fig. S5B**). To address this potential confounding factor further, we calculated the enrichment of crosstalk interactions in genes nominated by GWAS for other neurological or neuropsychiatric traits (**table S5**). We only used crosstalk interactions predicted from the control donors from this study to make results comparable across traits. This approach is an orthogonal strategy to determine if the abundance of microglia genes in the CellPhoneDB database skewed our previous enrichment results. Our reasoning was that if the crosstalk interactions observed in this study were biased towards microglia or other cell types due to database overrepresentation, we would expect to observe skewed enrichment patterns across traits. On the contrary, we observed distinct crosstalk enrichment patterns across neuropsychiatric traits (**Fig. 1G**). For example, crosstalk interactions involving OPCs were significantly associated with genes from one schizophrenia GWAS (association = 1.00, adj. *p* = 0.007). We also observed a nominally significant association for inhibitory neurons in genes identified in one major depressive disorder (MDD) GWAS (association = 0.57, *p* = 0.02). Importantly, we replicated the enrichment for microglia crosstalk interactions in genes from two AD GWAS (Jansen *et al.* and Marioni *et al.* Association = 1.16 and 1.05, adj. *p* = 0.016 and 0.006, respectively) *(23, 24)*. These results confirm that the crosstalk enrichment patterns across cell types are specific to each trait, and we did not observe a skew towards microglia or endothelial cells in any of the other traits despite these two cell types expressing more genes participating in CellPhoneDB interactions. This approach also independently highlights the crucial role of microglia mediating AD genetic risk.

Finally, we addressed the potential for bias resulting from the selection of candidate AD genes for our crosstalk analyses. The complex task of identifying causal genes in AD GWAS can hinder the accurate determination of cell types mediating AD genetic risk at individual loci. In addition, the possibility of nominating multiple genes within the same locus, likely participating in similar pathways (*e.g.,* the MS4A locus *(25)*), could lead to over-representation (“double-counting”) of the same GWAS signal. To mitigate these biases, we adopted a data-driven approach for nominating AD genes, relying strictly on cell type-specific chromatin co-accessibility between gene promoter regions and a fine-mapped AD GWAS variant *(12)* or the direct overlap of fine-mapped variants at the gene promoter region (Methods). This stringent approach nominates candidate AD genes and their corresponding cell types solely based on direct evidence from a public brain snATAC-seq dataset *(17)*. Despite microglia being among the least abundant cell types in the snATAC-seq dataset analyzed, we observed a two-fold higher enrichment for microglia crosstalk interactions in AD compared to using our original AD genes list (association = 1.51, *p* = 1.89e-4; **Fig. 1H**). This result shows that the enrichment of AD-related crosstalk interactions is robust to varying degrees of stringency in the strategy for selecting candidate AD genes and independently recapitulates the well-established role of microglia in mediating AD genetic risk.

Combined, these results indicate that the observed crosstalk enrichment patterns were highly robust to potential technical confounding factors. Furthermore, these analyses highlight that our crosstalk framework is highly flexible and can be extended to understand biological processes associated with other neurological and neuropsychiatric diseases.

### Microglia and neurons crosstalk interactions regulate additional known AD genes in microglia

Given our previous results prioritizing neuron-microglia crosstalk interactions in AD, we sought to investigate how the crosstalk signals between neurons and microglia could regulate gene regulatory networks downstream in microglia. Using a systems biology approach based on extending the functionality of the CytoTalk software *(26)* (Methods), we reconstructed the gene co-expression networks upstream of the crosstalk ligands and downstream of the receptors. CytoTalk is complementary to CellPhoneDB, as the latter does not inform the biological processes likely downstream of crosstalk interactions. In addition, the crosstalk interactions prioritized by Cytotalk reflect their predicted regulatory impact based on the co-expression network topology. Importantly, we did not restrict the crosstalk interactions in CytoTalk to those involving AD genes to allow an unbiased crosstalk prioritization. This way, any AD-related crosstalk interactions prioritized by Cytotalk reflect their predicted importance in modulating central genes in their respective co-expression networks.

Using CytoTalk, we reconstructed the gene regulatory network associated with crosstalk interactions between excitatory neurons and microglia for each donor category, which was then combined into a single network to help understand the broader biological processes likely regulated by neuron-microglia crosstalk (**Fig. 2A-B**). We focused on excitatory neurons because they had the highest association of AD-related crosstalk interactions with microglia and were the most represented broad neuronal subtype in our data (**Fig. 1A** and **1F**). The microglia crosstalk network identified by CytoTalk was enriched for immune processes, including phagocytosis and cytokine production (**Fig. 2C**, consistent with neuron-microglia crosstalk interactions modulating microglia activation states *(27)*. Strikingly, the microglia co-expression network downstream of the prioritized neuron-microglia crosstalk interactions was enriched for genes previously associated with AD, even after statistically accounting for the overrepresentation of AD-related genes expressed in microglia (Methods; cases OR = 3.50, adj. *p* = 3.92e-5; **Fig. 2D**). These results suggest that neuron-microglia crosstalk interactions propagate signals that modulate genes previously implicated in AD and involved in regulating microglial activation.

**Fig. 2:**
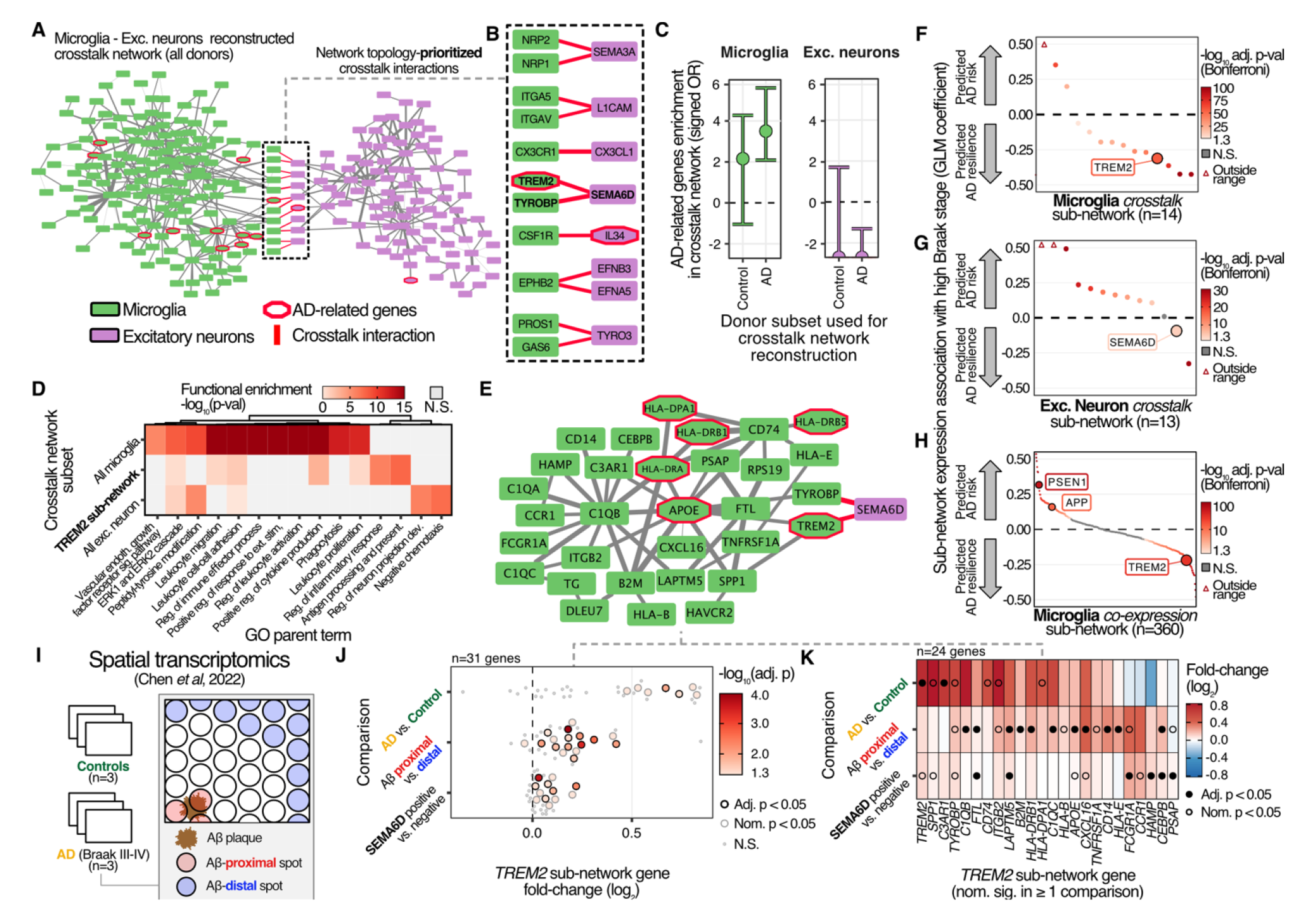
Crosstalk interactions between neurons and microglia are predicted to modulate AD risk genes. **A)** Microglia-excitatory neuron crosstalk network inferred by CytoTalk. **B)** Prioritized crosstalk interactions using CytoTalk. **C)** Enrichment of AD genes in the microglia crosstalk network across donor categories. **D)** GO enrichments for the genes participating in microglia and excitatory neurons crosstalk networks. **E)** Predicted TREM2 crosstalk subnetwork **F)** Association of microglia crosstalk subnetworks with high Braak stages. **G)** Association of all microglia subnetworks with Braak stages. **H)** Association excitatory neuron crosstalk subnetworks with high Braak stage. **I)** Spatial transcriptomics validation cohort overview. **J)** Changes in gene expression in the *TREM2* crosstalk sub-network associated with disease status, Aβ plaque proximity, and presence of *SEMA6D*-expressing cells. **K)** Individual gene view for comparisons in (J). GLM: generalized linear model.

Among the seven crosstalk interactions prioritized by CytoTalk based on the co-expression network topology, we identified the interaction between the neuronal ligand semaphorin 6D (SEMA6D) and TREM2/TYROBP (DAP12) (**Fig. 2B)**. This crosstalk interaction was initially described in the context of peripheral myeloid cells activation *(28)*, but its role in microglia and AD remains unknown. Given the central role of TREM2 in AD genetic risk, this notable knowledge gap motivated us to pursue this interaction further.

The TREM2-SEMA6D crosstalk is mediated by PLXNA1 *(28)*. Because of the low detection rate and limited dynamic range of *PLXNA1* in our snRNA-seq data (only ∼10% of microglia had detectable *PLXNA1* levels; max. *PLXNA1* expression = 3 reads; **fig. S6**), *PLXNA1* was not included in the CytoTalk reconstructed network. This is a reported limitation of snRNA-seq for lowly expressed genes *(29, 30)* and precluded the reconstruction of the *PLXNA1* co-expression network by CytoTalk, resulting in a direct link between *SEMA6D* and *TREM2/TYROBP* in the excitatory neuron-microglia network. Nonetheless, microglial *PLXNA1* and neuronal *SEMA6D* expression patterns were sufficiently specific for both CellPhoneDB and CytoTalk independently detect and prioritize the SEMA6D-PLXNA1/TREM2 crosstalk interaction between microglia and neurons in our analyses.

### The SEMA6D-TREM2 crosstalk axis is predicted to modulate microglia activation

We next sought to understand how the TREM2-SEMA6D crosstalk interaction could regulate microglia biology. We identified a sub-network comprised of genes highly connected to *TREM2* and *TYROBP* by partitioning the microglia crosstalk network into sub-networks (Methods). Our crosstalk network reconstruction analysis predicted that this *TREM2* sub-network is the target of neuronal SEMA6D (**Fig. 2E**). Furthermore, the *TREM2* sub-network was enriched for microglia activation pathways, indicating that we recapitulated the well-established link between TREM2 and microglia activation *(31)* through this unsupervised approach (**Fig. 2D**). Interestingly, in addition to genes linked to microglia activation, the TREM2 crosstalk sub-network included *APOE* and *HLA* genes, previously reported as AD risk genes. The co-expression of *TREM2* and *APOE* is consistent with studies showing that APOE is a TREM2 ligand *(32, 33)*. We identified a similar sub-network at the interface of the reconstructed inhibitory neurons and microglia crosstalk network, indicating that this is a general feature of neuron-microglia communication (**fig. S7A**). These results suggest that the TREM2-SEMA6D crosstalk interaction modulates AD risk genes in microglia and is a core feature of neuron-microglia communication.

To validate these results, we repeated the CytoTalk analyses in the snRNA-seq studies from the SWDBB, Rush ADRC, and UCI MIND ADRC cohorts *(16, 17, 22)*. Consistent with our results, CytoTalk prioritized the SEMA6D-TREM2 signaling axis mediating the crosstalk interactions between excitatory neurons and microglia in all three cohorts, as well as identifying a similar *TREM2* subnetwork in all but one of the datasets (**fig. S7B**). Finally, we determined that the *TREM2* sub-network and its predicted modulation by SEMA6D were robust to the choice of donors and the number of nuclei used to reconstruct the crosstalk network (**fig. S7C**). Together, these results reinforce that the unsupervised methodological approach in this study identified core elements of microglia gene regulation, which are predicted to be modulated by neuron-microglia cellular crosstalk interactions.

### The microglia *TREM2* sub-network expression is negatively associated with late AD stages

We next sought to understand how the *TREM2* sub-network related to AD progression. We leveraged the wide range of neuropathological states in our dataset to develop a statistical framework to test the association of this sub-network gene expression level with disease severity while controlling for genetic and other confounding factors (Methods). By analyzing gene expression at the level of gene sub-networks, this approach also helped mitigate data sparsity in snRNA-seq differential expression analysis. Given the comprehensiveness of Braak staging among the neuropathological annotations within our cohort, we used high Braak stage (Braak ≥ IV) as a surrogate of AD severity. Using this approach, we determined that the expression of the *TREM2* sub-network was negatively associated with high Braak stage (beta = −0.31, adj. *p* = 4.32e-57), indicating that this sub-network is downregulated in later AD stages. To understand this result within the broader context of all neuron-microglia crosstalk interactions, we calculated the association of all neuron-microglia crosstalk sub-networks with high Braak stage. Remarkably, the majority (11 of 14) of the microglia crosstalk sub-networks were negatively associated with high Braak stage, and the *TREM2* sub-network was among the most negatively associated with high Braak stage (**Fig. 2F**). These results suggest that neuron-microglia crosstalk interactions and their downstream targets in microglia are impaired in the later stages of AD.

Next, we sought to determine if SEMA6D is a potential modulator of the *TREM2* microglia crosstalk sub-network. We reasoned that if this were the case, the neuronal *SEMA6D* crosstalk sub-network association with Braak stage would agree in direction with the *TREM2* sub-network. Indeed, the neuronal *SEMA6D* sub-network was also negatively associated with high Braak stage (beta = −0.09, adj. *p* = 1.63e-04; **Fig. 2G**). Therefore, our findings indicate that the biological processes associated with the SEMA6D-TREM2 neuron-microglia crosstalk interactions are disrupted in AD and likely play a protective role by regulating TREM2- dependent microglia activation.

### Multiple microglia co-expression sub-networks are disrupted during AD progression

To gain further insights into the role of microglia in AD, we next adapted the network analysis framework of CytoTalk to analyze the transcriptome-wide microglia co-expression network (*i.e.*, using all expressed genes in microglia instead of the subset prioritized by CytoTalk; Methods). This approach allowed us to test the association of all microglia sub-networks with high Braak stage, regardless of the presence of reported ligands/receptors in the sub-networks. We partitioned this broader reconstructed microglia co-expression network in 360 sub-networks and independently recapitulated several sub-networks from the previous crosstalk-prioritized network reconstruction, including the *TREM2* sub-network (**fig. S8**). These microglia sub-networks were divided between positive and negative associations with high Braak stage (**Fig. 2H; table S6**). Consistent with the well-established roles of PSEN1 and APP in AD onset *(34)*, the *PSEN1* and *APP* co-expression sub-networks were among the most positively correlated with high Braak stage (betas = 0.32 and 0.16, adj. *p* = 8.25e-57 and 2.17e-13, respectively). In contrast, the sub-network of *SORL1*, a gene associated with protective roles in AD *(35, 36)*, was negatively associated with Braak stage (beta = −0.15, adj. *p* = 3.52e-11). Interestingly, our unsupervised approach identified two separate sub-networks with opposing directions of effect for the genes in the MS4A locus, which genetically controls soluble TREM2 levels *(25)* (*MS4A4A* and *MS4A6A* betas = −0.11 and 0.19, adj. *p* = 2.87e-06 and 2.93e-20, respectively). This result suggests that the MS4A genetic signal regulates at least two independent biological processes, consistent with what we reported in a previous study *(25)*. Notably, within the context of all microglia genes, the *TREM2* sub-network was among the most negatively associated with Braak stage (**Fig. 2H**), consistent with our analysis of the crosstalk-prioritized network. These results indicate that multiple biological pathways downstream of microglia-neuronal crosstalk are disrupted in AD. Furthermore, our unsupervised computational framework identified impaired TREM2-dependent microglia activation associated with AD progression.

### The TREM2 sub-network expression correlates with proximity to A**β** plaques and is up-regulated in the presence of SEMA6D

Our previous findings that the *TREM2* sub-network was among the most negatively associated with advanced AD stages motivated us to better understand how its expression changed as a function of neuropathological burden. To do so, we reanalyzed spatial transcriptomics profiles (10X Genomics Visium) from three control and three AD (Braak III and IV) human brains *(37)* and quantified the effects of local neuropathology, in particular proximity to Aβ plaques, in relation to gene expression patterns (**Fig. 2I**).

We first analyzed the global changes in gene expression between AD cases and controls and identified only 7 of 31 genes in the *TREM2* sub-network with nominal significant association (median log_2_ fold-change = 0.67; **Fig. 2J-K)**. However, when we compared Aβ plaque-proximal to Aβ plaque-distal regions, we observed significant up-regulation of most genes in the *TREM2* sub-network (17 of 31 genes at least nominally significant, median log_2_ fold-change = 0.18). The spatially resolved data also showed a progressive overexpression of genes in the *TREM2* sub-network as a function of Aβ plaque proximity (**fig. S9A-B**), indicating that this pathway is likely involved in the immune response to amyloid pathology. Supporting this hypothesis, we observed that other gene signatures linked to plaque-associated microglia in single-cell transcriptomics studies of AD mouse models *(15, 38)* were also overexpressed with Aβ proximity (**fig. S9C**).

Lastly, we leveraged the resolved spatial relationship of this dataset to test whether the *TREM2* sub-network expression levels changed in proximity to *SEMA6D*-expressing cells. In line with our single-cell analyses, we observed a significant up-regulation of the *TREM2* sub-network when comparing *SEMA6D*-positive versus *SEMA6D*-negative Visium spots (19 of 31 genes at least nominally significant; median log_2_ fold-change = 0.081; **Fig. 2J-K**). The level of *TREM2* sub-network activation in proximity to *SEMA6D* was comparable between cases and controls (**fig. S9D**). These results, combined with the lower expression of the *TREM2* subnetwork in the high Braak stages donors from the snRNA-seq data (**Fig. 2H**), suggest that the *TREM2* crosstalk sub-network is active during earlier Braak stages, responds to local neuropathology (Aβ plaques) and SEMA6D signaling, but loses function as the disease progresses. Our findings suggest that the *TREM2* sub-network is involved in the response to Aβ plaques and is activated by SEMA6D.

### SEMA6D induces immune activation in iPSC-derived microglia in a TREM2-dependent manner

To elucidate the role of SEMA6D-TREM2 crosstalk in microglia function, we used a human iPSC-derived microglia model *(39)* (iMGL; **fig. S10A**; Methods) that expresses established microglia markers, including TREM2, IBA1, and TMEM119 (**fig. S10B**). In addition, we generated *TREM2* KO human iPSCs using CRISPR/Cas9 to examine the role of the SEMA6D-TREM2 signaling axis on microglia function (**fig. S10C**). We verified the loss of TREM2 expression at the protein level in the KO cell line by western blot analysis (**fig. S10D**).

Because microglia regulate brain homeostasis through phagocytic activity and modulate neuroinflammation by releasing immune cytokines *(40–42)*, we performed phagocytosis and cytokine assays (**fig. S11**). To determine if SEMA6D can regulate iMGL phagocytic activity and whether this process is TREM2 dependent, we treated WT and *TREM2* KO iMGL with recombinant SEMA6D protein. We measured the degree of phagocytic activity using pHrodo-labeled human synaptosomes as the phagocytic cargo. We observed increased phagocytosis in WT iMGL treated with SEMA6D starting at 6 hours of treatment with SEMA6D (1.3-fold change increase at 24 hours, *p* = 1.50e-8). In contrast, *TREM2* KO iMGL treated with SEMA6D had a less pronounced increase in phagocytosis compared to untreated *TREM2* KO cells (1.1-fold change increase at 24 hours, *p* = 0.001; **Fig. 3A-B**; **fig. S12A)**. In parallel, we analyzed conditioned media of WT and *TREM2* KO iMGL using a multiplex immunoassay to determine if SEMA6D can regulate iMGL cytokine release. We found that SEMA6D increased the secretion of TNF-α by and IL-6 in WT but not *TREM2* KO iMGL (WT TNF-α and IL-6 fold-changes = 1.37 and 3.59, *p* = 4.58e-5 and 3.11e-5, respectively; **Fig. 3C**). We replicated the effects of SEMA6D treatment in iMGL generated from an independent WT isogenic iPSC line, indicating that the observed effects of SEMA6D treatment in iMGL activation were not due to cell line-specific effects (**fig. S12B-C**). Together, these results indicate that SEMA6D increases iMGL phagocytosis and secretion of TNF-α and IL-6 cytokines in a primarily TREM2-dependent manner.

**Fig. 3.**
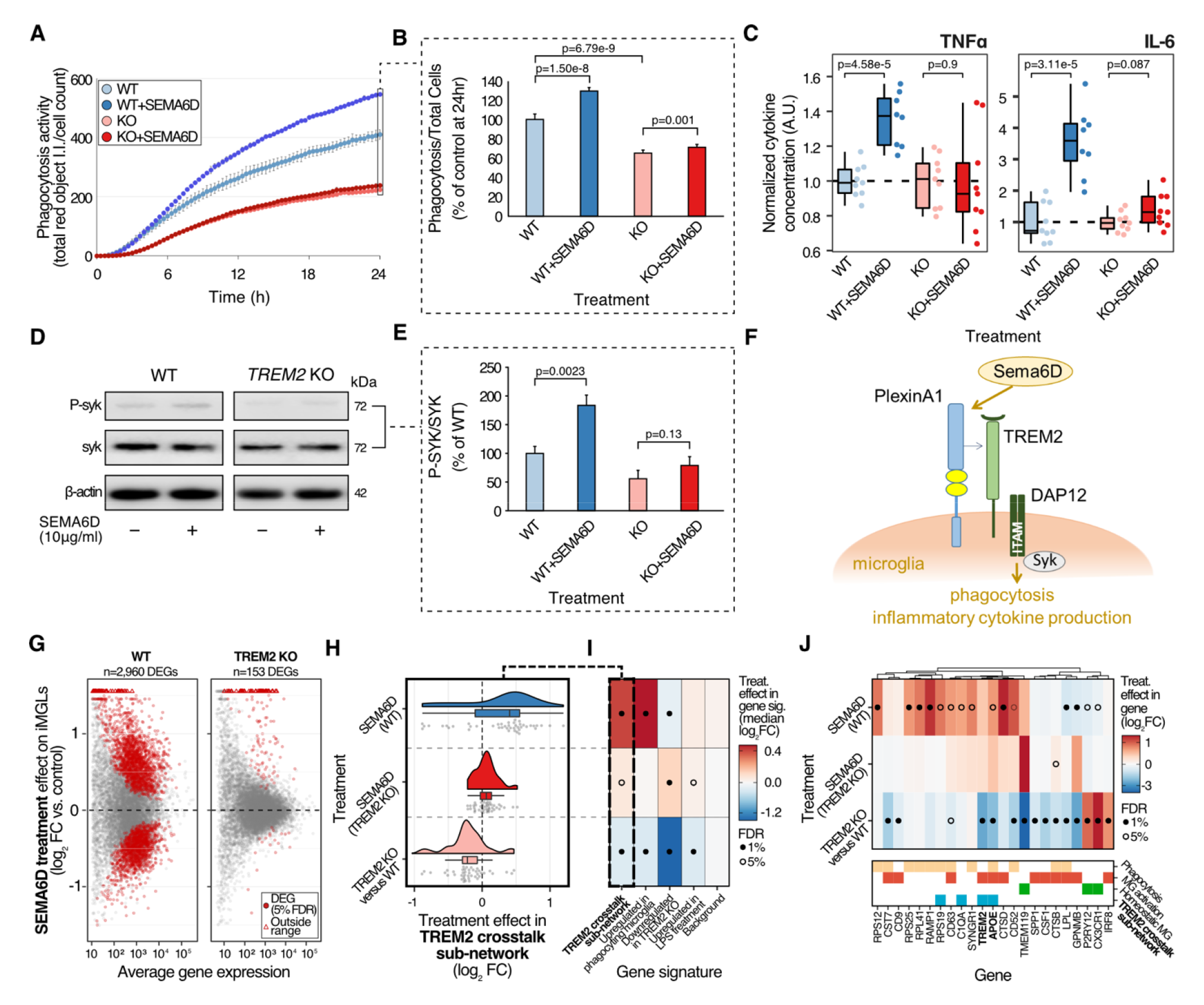
SEMA6D treatment induces microglia activation in a TREM2-dependent manner. **A)** Phagocytosis of synaptosomes by WT or *TREM2* KO iMGL treated with SEMA6D (5 μg/ml). I.I., integrated intensity. Data averaged from three independent experiments (shown in fig. S12); mean ± SD values. **B)** Quantification of phagocytosis of synaptosomes by WT or *TREM2* KO iMGL treated with SEMA6D at 24 hr. Data represent three independent experiments performed in triplicate; mean ± SEM values at 24 hrs, *p*-values obtained from meta-analyzing the three experiments. **C)** Quantification of media from WT or *TREM2* KO iMGL treated with SEMA6D (10 μg/ml) for cytokine release of TNF-α and IL-6, using the Mesoscale V-plex neuroinflammation panel. Boxplots indicate median and interquartile ranges, *p*-values calculated using linear regression. **D)** Representative western blot analysis of p-SYK and total SYK in WT and *TREM2* KO iMGL treated with SEMA6D (10 μg/ml). β-actin as loading control **E)** Quantification of p-SYK/total SYK western blot analysis, n=3 (D). **F)** Schematic of the proposed SEMA6D-TREM2/DAP12 signaling complex. **G)** RNA-seq effect size distribution on SEMA6D-treated iMGL (WT and *TREM2* KO). **H)** Transcriptional changes in the *TREM2* crosstalk sub-network associated with SEMA6D treatment (WT or *TREM2* KO vs. untreated control) or *TREM2* KO (*TREM2* KO vs. WT, no treatment). Only *TREM2* sub-network genes differentially expressed in at least one comparison were included. **I)** Transcriptional effects of the same conditions as (B) across biologically relevant gene signatures. Background corresponds to 500 randomly selected genes. Solid dots correspond to a 1% FDR significance threshold for comparing the effect size distribution to the corresponding background (Mann-Whitney test). **J)** Transcriptional effects of the experimental conditions across a representative subset of highly differentially expressed genes from the signatures in (I).

TREM2 mediates signaling through the adaptor protein TYROBP (DAP12), and the activation of TREM2 results in tyrosine phosphorylation within the ITAM motif and subsequent SYK phosphorylation *(43)*. To determine if SEMA6D activates TREM2 downstream signaling, we analyzed WT and *TREM2* KO iMGL protein lysates for phosphorylated SYK expression (p-SYK) normalized to total SYK expression (SYK). Treatment of WT iMGL with SEMA6D induced a 1.84-fold increase in SYK phosphorylation (*p* = 0.0023), but these effects were not significant in *TREM2* KO iMGL (*p* = 0.13; **Fig. 3D-E**; **fig. S13**). These results demonstrate that SEMA6D can directly activate TREM2 signaling and suggests that SEMA6D signals through the TREM2/TYROBP (DAP12) complex in microglia (**Fig. 3F**), although we cannot exclude that SEMA6D simultaneously activates other signaling pathways.

To systematically characterize the transcriptional changes induced by SEMA6D treatment in iMGL, we generated bulk RNA-seq data for the SEMA6D-treated WT and *TREM2* KO iMGL and the corresponding untreated controls (**fig. S13**). We observed significant transcriptional changes associated with *TREM2* KO (n = 1,408 differentially expressed genes at 5% FDR; **fig. S13A-B**). As expected, *TREM2* was among the most downregulated genes in the *TREM2* KO iMGL (adj. *p* = 4.30e-22, rank = 26; **fig. S13A)**. Strikingly, we observed robust transcriptional changes in SEMA6D-treated WT iMGL but not in SEMA6D-treated *TREM2* KO iMGL (n = 2,960 and 153 differentially expressed genes at 5% FDR, respectively; **Fig. 3G; fig. S13A-B**), consistent with a pivotal role for TREM2 in mediating SEMA6D signaling in microglia. To further understand how TREM2 mediates this signaling pathway, we focused on the *TREM2* co-expression crosstalk sub-network predicted from the snRNA-seq data (**Fig. 2E**). The *TREM2* sub-network had significantly lower expression in the untreated *TREM2* KO iMGL than WT (median log_2_ fold-change = −0.22, adj. *p* = 4.76e-04), consistent with TREM2 being a key regulator of this sub-network. In line with this interpretation, the *TREM2* sub-network was activated by SEMA6D treatment in the WT iMGL (median log_2_ fold-change = 0.40, adj. *p* = 7.50e-3) but significantly less so in the *TREM2* KO iMGL (median log_2_ fold-change = 0.06, adj. *p* = 0.040; **Fig. 3H-I**). These results are consistent with the TREM2 signaling pathway being the primary mediator of SEMA6D in microglia but also suggest that SEMA6D can activate other pathways in microglia to a lesser extent.

We next analyzed biologically relevant transcriptional signatures previously described in microglia to gain further insights into how SEMA6D treatment regulates microglial transcriptional programs. These included genes up-regulated in phagocytosing microglia *(41)* and those up-regulated in response to LPS treatment *(18)*. As control signatures, we included genes down-regulated in another *TREM2* KO iMGL dataset *(44)* and a set of randomly selected genes for which we would not expect any concerted transcriptional changes. We observed the strongest effects of SEMA6D treatment in the phagocytosing microglia gene signature (median log_2_ fold-change = 0.47; **Fig. 3I**), indicating that SEMA6D activates genes involved in phagocytosis in the WT but not *TREM2* KO iMGL. Notably, *TREM2*, *APOE*, and *RPS19* are among the most up-regulated genes by SEMA6D treatment in WT iMGL. These genes are either present in the phagocytosing microglia gene signature or correspond to genes previously linked to microglia activation *(38, 15)* (**Fig. 3J**). Our results indicate that SEMA6D-TREM2 crosstalk signaling induces a TREM2-mediated cascade of transcriptional events resulting in microglia activation.

## DISCUSSION

In this study, we leveraged single-nucleus gene expression profiles from a diverse cohort of brain donors to systematically dissect the contribution of cross-cellular signaling (cellular crosstalk) networks to AD. Our data-driven approach to identifying active crosstalk interactions and reconstructing their corresponding downstream pathways provides additional evidence that disrupted cellular crosstalk networks contribute to neurodegeneration. One remarkable finding from our study is that a significant portion of AD risk genes is either directly involved in crosstalk interactions or immediately downstream of crosstalk interactions involving microglia.

These results highlight the difficulty of characterizing the prominent role of microglia in AD, as the integration of complex signals originating in other brain cell types is core to their function. Specifically, our results support that dysregulation of the intricate signaling between neurons and microglia is linked to AD progression. Therefore, focusing on cellular crosstalk networks can provide further functional context to understand the biology of genes associated with AD risk in a cell-autonomous and non-autonomous manner.

Among the interactions we detected between neurons and microglia, we identified a functional link between neuronal SEMA6D and microglial TREM2. Semaphorins and their receptors regulate immune cell function and are genetically and functionally implicated in AD *(6, 45–47)*. In the brain, semaphorin signaling was initially described as a mediator of axon guidance via the plexin family of receptors *(48)*. However, a growing body of evidence indicates these molecules are involved in immune responses *(5, 28, 45, 49–51)*. The role of SEMA6D in immune activation was described in a study showing that SEMA6D induces activation of bone marrow-derived macrophages in a TREM2- and PLXNA1-dependent manner via the activation of DAP12, consistent with the formation of a complex *(28, 52, 53)*. Additional studies also linked semaphorin signaling to immune activation and neurodegeneration *(5, 51, 54–56)*. However, despite the role of SEMA6D in TREM2-dependent immune activation of peripheral myeloid cells being described over a decade ago (which allowed us to computationally test this interaction in the first place) and the well-established role of TREM2 in AD genetic risk, there is a notable gap of understanding regarding this signaling pathway in the context of microglia and AD. By leveraging iMGL, we demonstrated that SEMA6D signaling induces a TREM2- dependent microglia activation phenotype marked by phagocytosis and inflammatory cytokine release and transcriptionally similar to phagocytosing microglia *(41)*. Nevertheless, it remains to be determined if SEMA6D induces a state of microglia that might be beneficial in clearing neuropathological changes associated with AD. Our observation that the *TREM2* co-expression sub-network is activated in the proximity of Aβ plaques and *SEMA6D*-expressing cells, combined with our observation that the transcriptional networks upstream and downstream of the SEMA6D-TREM2 interaction are downregulated in late AD stages, suggest that loss of this interaction exacerbates the deleterious processes occurring in the later stages of this disease.

Our single-cell transcriptomics analyses implicated excitatory neurons as the primary partners for microglia regarding the SEMA6D-TREM2 crosstalk interaction. However, we also observed *SEMA6D* expression in other neuronal subtypes and, to a lesser extent, in other cell types. Therefore, further experimental studies, such as cell co-cultures, are necessary to precisely determine the primary cell types contributing to SEMA6D-TREM2 signaling in microglia.

A previous study showed that SEMA6D promotes peripheral dendritic cell activation and osteoclast differentiation via the receptor complex harboring PLXNA1 and TREM2 *(28)*. Thus, it is conceivable that SEMA6D functions as a natural ligand for the PLXNA1/TREM2 co-receptor and enhances TREM2 signaling in human microglia. Therefore, neuronal SEMA6D could influence functional properties and fate via the stimulation of TREM2-dependent intracellular signaling and induction of the *TREM2* gene expression network. However, other studies described how SEMA6D also regulates lipid metabolism and polarization of macrophages via the interaction with another class A plexin family member, Plexin A4 *(49, 57)*. Plexin A4 coding variants have been linked to AD risk *(49, 58)* and found to modulate amyloid and tau pathology *(54)*. Thus, semaphorin-plexin signaling may play a fundamental role in regulating the functional interactions with microglia and other cell types and may be perturbed in AD. Given that we observed weak transcriptional effects in iMGL associated with SEMA6D treatment in the absence of *TREM2*, it is possible that other proteins, such as Plexin A4, act as secondary SEMA6D receptors in microglia. Therefore, future studies are necessary to determine the complete network of proteins mediating SEMA6D crosstalk in microglia.

While this study focused on a restricted subset of crosstalk interactions involving microglia and neurons, our systematic characterization of cross-cellular signaling patterns identified thousands of candidate interactions involving all brain cell types represented in our snRNA-seq data. Several of these interactions warrant further investigation. For example, the interleukin receptor IL1RAP has been previously implicated in genetic studies of AD endophenotypes *(3, 59–61)*, and the contribution of IL-1 signaling to neurodegenerative diseases is well-established *(62–64)*. In line with these studies, we identified the *IL1RAP* sub-network in microglia as the most negatively associated with Braak stage (**table S6**). These results suggest that the IL-1 signaling pathway disruption is likely involved in AD progression. More broadly, the IL1RAP case highlights that the continued exploration of brain crosstalk networks identified in this study will yield valuable biological insights into AD biology.

A limitation of our study is its reliance on existing databases of curated crosstalk interactions, which exclude interactions not yet reported in the literature. Moreover, our transcriptomics-based approach may overlook cellular communication mediated by molecules synthesized through complex biochemical pathways lacking canonical ligand genes (*e.g.*, lipids and some neurotransmitters) or not relying on a specific receptor in the conventional sense (*e.g.*, nitric oxide signaling). Finally, our understanding of the genetic risk of AD and other neuropsychiatric traits is incomplete. This knowledge gap hinders not only the discovery of yet-unknown risk genes but also their corresponding crosstalk networks, thus precluding a complete characterization of the role of cellular crosstalk in neurodegeneration. Despite these constraints, our results indicate that a systematic characterization of cellular crosstalk networks can provide valuable insights into the biology of neurodegenerative diseases, potentially aiding in identifying novel therapeutic targets. Given our findings, we advocate for developing new high-throughput assays to systematically identify cell-to-cell communication pathways.

Finally, we identified unique crosstalk enrichment patterns for genes found in genetic studies of other neurological or neuropsychiatric traits. This underscores the integral role of cellular crosstalk in normal brain physiology and suggests that acknowledging this regulatory layer could aid in understanding how candidate disease risk genes fit into biological pathways. Together, our findings strongly support that the systematic characterization of cellular crosstalk networks is a viable strategy for gaining insight into the biology of neurodegenerative diseases and nominating targets for novel therapies.

## MATERIALS AND METHODS

### Study design

Samples were previously obtained with informed consent for research use and were approved by the review board of Washington University in St. Louis. AD neuropathological changes were assessed according to the National Institute on Aging-Alzheimer’s Association (NIA-AA) criteria. Demographic, clinical severity and neuropathological information are available in our original study *(14)*.

### iMGL experiments

All experimental procedures are described in the Supplementary Materials of this manuscript.

### Statistical analyses

Detailed computational and statistical methods are described in the Supplementary Materials of this manuscript.

## Data and materials availability

Human iMGL bulk RNA-seq data are available under GEO accession GSE226507. The snRNA-seq data from the Knight ADRC is publicly available by request from the National Institute on Aging Genetics of Alzheimer’s Disease Data Storage Site (NIAGADS) under accession number NG00108 (https://www.niagads.org/datasets/ng00108). DIAN brain bank snRNA-seq data access requires a request through https://dian.wustl.edu/our-research/for-investigators. All scripts necessary to reproduce the figures from this manuscript will be available at https://github.com/albanus-research/2021_trem2_adad_snRNAseq. The modified CytoTalk version used for this manuscript is available at https://github.com/rdalbanus/CytoTalk.

## Acknowledgments

This manuscript has been reviewed by DIAN Study investigators for scientific content and consistency of data interpretation with previous DIAN Study publications. We acknowledge the altruism of the participants and their families and the contributions of the DIAN research and support staff at each participating site for their contributions to this study. The data available in the AD Knowledge Portal would not be possible without the participation of research volunteers and the contribution of data by collaborating researchers. The results published here are in whole or in part based on data obtained from the AD Knowledge Portal (https://adknowledgeportal.org). This work was supported by access to equipment made possible by the Hope Center for Neurological Disorders and the Departments of Neurology and Psychiatry at Washington University School of Medicine. We thank Barbara Corneo for advice on iPSC protocols and Wei Wang and Caisheng (Luke) Lu, MD, Ph.D., for assisting with FACS sorting. iMGL cell sequencing and preliminary RNA-seq data processing were conducted by Novogene.

## Author contributions

RDA analyzed data, designed and performed computational experiments, wrote the manuscript. GMF analyzed data, designed and performed iPSC experiments, wrote the manuscript. LB performed initial snRNA-seq quality control, cell clustering, and integration with public datasets. SC annotated spatial transcriptomics data. QG performed spatial transcriptomics deconvolution. AK designed and performed experiments, analyzed data, and revised the manuscript. MA performed experiments and revised the manuscript. SFY performed computational experiments. BCN performed initial snRNA-seq processing quality control. PMRP revised the manuscript. DMH revised the manuscript. JCM supervised sample acquisition and revised manuscript. EMD supervised sample acquisition, revised manuscript. MF supervised sample acquisition and revised the manuscript. JPC supervised sample acquisition, revised manuscript. RJP supervised and participated in sample acquisition, performed neuropathological assessments, and revised the manuscript. EEM contributed to experimental data acquisition. BAB analyzed data, supervised snRNA-seq data generation, and revised the manuscript. LP conceptualized results and revised the manuscript. GTS supervised sample and genetic data acquisition, revised manuscript. QM supervised spatial transcriptomics computational analyses. HF: Supervised sample acquisition and revised manuscript. CMK supervised experimental data acquisition and revised the manuscript. OH conceptualized and designed computational experiments, analyzed data, supervised and edited the manuscript, and supervised all aspects of the project. TWK conceptualized and designed iPSC experiments, analyzed data, supervised and edited the manuscript, and supervised all aspects of the project.

## Supplementary Materials

Materials and Methods

Fig. S1 to S14

References (65-85)

### Other Supplementary Material for this manuscript includes the following

Table S1 to S6 (Excel files)

