## Supplementary Material for "Systematic analysis of cellular crosstalk reveals a role for SEMA6D-TREM2 regulating microglial function in Alzheimer’s disease"

### MATERIALS AND METHODS

#### snRNA-seq data processing

We reanalyzed single-nucleus transcriptomic profiles (snRNA-seq) of superior parietal cortex tissue samples from 67 donors of the Knight Alzheimer Disease Research Center (Knight ADRC) and the Dominantly Inherited Alzheimer Network (DIAN) *(14)*. Briefly, we used the 10X Chromium single-cell Reagent Kit v3, aiming for 10,000 nuclei per individual and 50,000 reads per nucleus, as recommended by 10X Genomics. We mapped the sequenced reads to the GRCh38 human reference genome using CellRanger (v. 6.1.1) and filtered nuclei based on sequencing depth and percent mitochondrial reads. We obtained snRNA-seq data from 294,114 nuclei after stringent QC *(14, 67)* (median nuclei per donor after QC = 3,637). We used Seurat *(68)* (v. 3.2.3) to identify cell types and transcriptional states using previously described cell type markers *(69)*.

We obtained the count matrices and barcode metadata from three other public snRNA-seq datasets from human brains *(16, 17, 22)* as made available by the authors. We used the existing barcode metadata from these studies to annotate the nuclei into cell types and donor groups (*e.g.*, AD, controls). We considered any nuclei not in the final metadata table for the respective study as failing QC. These nuclei were discarded from downstream analyses. In order to run CellPhoneDB and CytoTalk in these datasets, we normalized each study separately using the SCT normalization *(70)* from Seurat.

#### Reanalysis of brain snATAC-seq data

We downloaded the raw snATAC-seq fastq files from the Morabito *et al.* study *(17)* and re-processed this dataset using CellRanger-ATAC (v. 2.0.0) count function. First, we filtered the resulting GRCh38 BAM files for each sample using flags *-f 3 -F 4 -F 8 -F 256 -F 1024 -F 2048 -q 30* to retain high-quality, properly mapped alignments. Second, we split each sample BAM file into separate files corresponding to each cell type using the barcode annotations reported by the authors. Then, for each cell type, we merged all BAM files across donors and called narrow peaks using MACS2 (v. 2.2.7.1) *(71)* using flags *--nomodel --shift -100 --extsize 200 --keep-dup all --call-summits -B*. Third, we filtered all peaks overlapping the GRCh38 ENCODE exclusion list regions (ENCODE accession ENCFF356LFX) *(72)*. Finally, we used CICERO *(73)*, with default parameters, to calculate co-accessibility across ATAC-seq narrow peaks for each cell type separately.

#### CellPhoneDB analyses

We used CellPhoneDB *(20)* (v. 2.1.7) to estimate crosstalk interactions based on the expression of known ligand-receptor pairs. We used the *cellphonedb method statistical_analysis* function with default parameters to calculate crosstalk interactions at the cluster level (transcriptional state) using the sctransform-normalized snRNA-seq count matrices as input. We performed CellPhoneDB analyses for each subset of donors separately (*e.g.*, controls, sporadic AD). We considered significant all those interactions that passed multiple testing correction (Bonferroni $p < 0.05$) for the entire set of analyses (all transcriptional states and donor categories).

To address potential bias from differences in cell type representation, we performed an additional CellPhoneDB analysis, downsampling each cluster in our snRNA-seq data to 100 nuclei barcodes. Similarly, we re-ran CellPhoneDB in multiple combinations of two to three cell types in our data (e.g., astrocytes and OPC only, astrocytes, microglia, and OPC only) to confirm that the resulting interactions were stable across subsets of cell types. Finally, we repeated the CellPhoneDB analyses individually for a subset of brains to confirm that the crosstalk patterns were identified at a single brain level. For other public snRNA-seq datasets analyzed in this study, we used the cluster and donor labels provided by the authors of each study.

To calculate the significance of CellPhoneDB interactions, we used Fisher exact tests (FETs) to compare the number of interactions at the cell type level. We generated one 2x2 contingency table per cell type and donor subset to compare controls versus AD. One dimension of the contingency table encoded the number of unique significant CellPhoneDB interactions involving the cell type of interest versus all other cell types combined. The other dimension encoded the number of interactions identified in the controls versus the donor category of interest. To calculate the enrichment of crosstalk interactions involving AD-related genes per cell type, we generated, for each donor category, 2x2 contingency tables encoding the number of interactions involving the cell type of interest (yes versus no) and involving AD genes (yes versus no). All FET p-values were corrected for multiple testing using the Bonferroni correction based on the total number of tests performed for each analysis.

To calculate the association of AD-related crosstalk interactions per cell type, we performed logistic regressions predicting whether the interaction involved an AD gene (binary variable) as a function of the cell types in which it was detected:

$$interaction\_is\_AD \sim cell\_type\_a + cell\_type\_b + (\ldots) + cell\_type\_n$$

To calculate the enrichment for cell-type pairs, we calculated the association of interactions with whether they were detected in a cell-type pair of interest versus not (*e.g.*, microglia and exc. neurons versus everything else), iterating across all possible cell-type pairs:

$$interaction\_is\_AD \sim detected\_in\_cell\_pair + not\_detected\_in\_cell\_pair$$

We performed these analyses for each dataset separately and in combination (mega-analysis). For the latter, we included a covariate one-hot-encoding each dataset:

$$interaction\_is\_AD \sim (covariates) + dataset\_A + dataset\_B + (\ldots) + dataset\_n$$

All logistic regression $p$-values were corrected for multiple testing using the Bonferroni correction based on the total number of cell types and datasets (if applicable).

**Crosstalk enrichments for other neuropsychiatric traits**

We downloaded the GWAS associations of neuropsychiatric traits from the European Bioinformatics Institute GWAS catalog (https://www.ebi.ac.uk/gwas) for the studies listed in **table S5**. We used the mapped genes for each locus as the input gene list to calculate the crosstalk enrichments in each study. We used the same FET approach described above for the AD-related crosstalk interactions to calculate the enrichment of genes nominated by each GWAS study (**Fig. 2D**). To calculate the enrichment of crosstalk interactions involving genes associated with AD GWAS based on co-accessibility (**Fig. 2E**), we used as the input gene list all genes with a transcription start site (TSS) region either 1) overlapping an ATAC-seq peak co-accessible with another ATAC-seq peak harboring a fine-mapped AD GWAS variant (PPA > 0.01) from the Schwartzentruber *et al.* study *(12)* or 2) overlapping an ATAC-seq peak in any cell type harboring a fine-mapped variant.

**Downstream crosstalk networks reconstruction using CytoTalk**

We used CytoTalk (v. 4.0.3) with minor modifications to identify the signaling networks downstream of crosstalk interactions. CytoTalk first reconstructs the co-expression network of two cell types using information theory, then prioritizes biologically relevant crosstalk interactions based on their connection to central genes in each network using graph theory. This analysis results in a prioritized network of genes predicted to mediate the crosstalk signals between the two cell types. Briefly, CytoTalk reconstructs the co-expression network for each cell type pair and connects them based on a database of ligand-receptor interactions. It then uses a prize-collecting Steiner forest algorithm to identify the crosstalk signaling network between the two cell types *(26)*. We generated input csv files per cell type for each donor category containing the sctransform-normalized snRNA-seq log-transformed counts per barcode. Additionally, we increased the stringency of the CytoTalk networks by manually inputting only interactions that were also present in CellPhoneDB. This step was done because CytoTalk uses protein-protein interactions and text mining from STRING-DB *(74)* to build its crosstalk interactions database, which we consider overly permissive. For the network visualization in **Fig. 2A**, we used the union of the crosstalk signaling network identified for each donor category in microglia versus excitatory neurons. We generated network visualizations using Cytoscape (v. 3.9.0) *(75)* and RCy3 (v. 2.2.0) *(76)* and identified crosstalk sub-networks using the community cluster (GLay) function of clusterMaker (v. 2.2) *(77)*. The modified CytoTalk version used in this study (https://github.com/rdalbanus/CytoTalk) fixed minor bugs and enabled better parallelization control for a cluster environment. These issues have since been fixed in newer CytoTalk releases.

To calculate the enrichment of AD-related genes downstream of the crosstalk interactions (**Fig. 2C**), we used FETs to compare the number of AD-related genes in each side of the excitatory neuron-microglia crosstalk network versus all AD-related genes expressed in the corresponding cell type. All FET p-values were corrected for multiple testing using the Bonferroni correction based on the total number of tests performed for each analysis.

**Network analyses**

We used the *compute_mutual_information_single* function from CytoTalk to reconstruct the excitatory neurons and microglia co-expression networks for each donor category. Briefly, this function calculates the mutual information (MI) *(78)* across all pairs of genes per cell type and then applies the ARACNE algorithm *(79)* to eliminate most indirect interactions between genes. We then merged the ARACNE-filtered MI matrices for each cell type across donor categories by keeping the highest MI value for each interaction. In order to identify co-expression sub-networks, we applied the WGCNA pipeline *(80)* on the merged MI matrix. To test the association of each sub-network (n = 360) with the Braak stage, we first calculated the eigengene (the first principal component of gene expression) of each sub-network. We then tested the association of each eigengene with a high Braak stage using the binomial regression model:

$$Braak\ high_i \sim eigengene_{ij} + sex_i + APOE4_i + TREM2_i$$

In this model, $i$ represents a barcode and $j$, a co-expression sub-network. $Braak\ high_i$ , $sex_i$, $APOE4_i$, and $TREM2_i$ are binary variables encoding the donor Braak stage (Braak ≥ IV versus Braak ≤ III), sex, *APOE* genotype (one or two copies of *APOE4* versus zero), and *TREM2* genotype (common variant versus R47H, R136W, or R62H). We ran this model for each sub-network and obtained the corresponding coefficient and p-value of the eigengene term. Then, we adjusted the significance for multiple testing using the Bonferroni correction. Similarly, we performed this same analysis in the sub-networks obtained from the CytoTalk microglia-excitatory neurons crosstalk network.

**Spatial transcriptomics**

We processed the raw count matrices from the 10X Genomics Visium data as described in the original study *(37)*. Briefly, we used only spots that passed QC in the original study and applied the SCT integration pipeline from Seurat to integrate data across donors. All differential expression analyses were performed using NEBULA *(81)*, using covariates for the donor of origin (random effects model) and the previously calculated proportion of neurons and microglia using Cell2location *(82)* for each Visium spot. All spots were previously manually annotated regarding the presence of Aβ plaques based on immunostaining from immediately adjacent tissue slices. For the comparisons between Aβ-proximal versus distal spots, we defined as Aβ-proximal spots any spot that directly overlapped Aβ plaques, and we defined as Aβ-distal spots any spot that was more than four spots away from the nearest Aβ-proximal spot. For the *SEMA6D*-positive versus -negative comparisons, we used a threshold of ≥1 SCT-normalized *SEMA6D* reads to separate between classes. For cases versus controls, we used all the spots for each donor.

**iMGL RNA-seq data analysis**

Human mRNA sequencing and bioinformatics analysis were performed by Novogene. Raw fastq files were mapped to the GRCh38 reference genome using HISAT2 (v. 2.0.5) *(83)* and the corresponding GRCh38 index files (http://daehwankimlab.github.io/hisat2/download). We used DESeq2 *(84)* for differential analyses. Briefly, we retained genes with ≥ 10 counts in at least three samples and processed all samples together to increase the statistical power to calculate dispersion. We then retrieved the fold changes and significances for the comparisons of interest individually (*e.g.*, SEMA6D-treated WT vs. untreated WT). Finally, we adjusted for multiple testing per comparison using the Benjamini-Hochberg correction *(85)* and used a 10% false discovery rate (FDR) threshold to identify differentially expressed genes.

**Functional enrichments**

All gene ontology (GO) functional enrichments in this study were calculated with the WebGestaltR R package (v. 0.4.4), using the *ORA* method, the *geneontology_Biological_Process_noRedundant* annotation, and the *genome_protein-coding* reference set. We used an FDR threshold of 1 for all analyses to recover all enrichments and manually adjusted p-values for multiple testing using the Bonferroni correction (significance threshold used: $p < 0.05$). We used the R package rrvgo (v. 1.9.1) to collapse redundancy in GO terms. For each WebGestaltR analysis, we generated a similarity matrix across all GO terms significant in at least one comparison using the *calculateSimMatrix* function. We then reduced the significant terms for each analysis using the *reduceSimMatrix* function. The *threshold* values for the *reduceSimMatrix* function were selected based on visual inspection of the resulting terms (0.8 for the analyses in Figs. 1F and 3C and 0.9 for Fig. 3F). Finally, we reported the most significant p-value among the grouped terms under each parent term.

**Cell culture**

Human iPSCs were maintained feeder-free on matrigel-coated plates in StemFlex medium (Gibco) in a humidified incubator (5% $CO_2$, 37 °C). Cells were passaged as aggregates using ReLeSR (StemCell Technologies), and media was changed daily.

**Generation of TREM2 KO iPSCs**

The WT iPSC line (APOE 3/3) was a generous gift from Dr. Barbara Corneo *(65)*. TREM2 KO iPSCs were generated using the Human Stem Cell Nucleofector™ Kit 2 (Lonza) according to the manufacturer's protocol. Briefly, iPSCs were resuspended in 100 µL nucleofection buffer and 5 µg of vector lentiCRISPRv2GFP expressing sgRNA against TREM2 (5'ATCACAGACGATACCCTGGG 3'). LentiCRISPRv2GFP was a generous gift from David Feldser (Addgene plasmid# 82416; http://n2t.net/addgene:82416; RRID:Addgene_82416). The suspension was transferred to the Amaxa Nucleofector cuvette and transfected using program CA-137. GFP-expressing cells were single-cell sorted into 96-well plates using the BD Influx cell sorter (Columbia University CCTI/HICCC Flow Core). Cell colonies were maintained and expanded. TREM2 KO cell lines were verified by Sequencing (Genewiz).

**Differentiation of iPSC to iMGL**

iPSC-microglia were generated as described in McQuade et al., 2018 *(39)*, using the iPSC lines described in the previous section. Briefly, iPSCs were directed down a hematopoietic lineage using the STEMdiff Hematopoietic kit (STEMCELL Technologies). After 12 days in culture, hematopoietic progenitor cells (HPCs) were transferred into a microglia differentiation medium containing DMEM/F12, 2× insulin-transferrin-selenite, 2× B27, 0.5× N2, 1× GlutaMAX, 1× nonessential amino acids, 400 μM monothioglycerol, and 5 μg/mL human insulin. Media was added to cultures every other day and supplemented with 100 ng/mL IL-34, 50 ng/mL TGF-β1, and 25 ng/mL M-CSF (PeproTech) for 28 days. In the final three days of differentiation, 100 ng/mL CD200 (Novoprotein) and 100 ng/mL CX3CL1 (PeproTech) were added to the culture. In addition to the WT and *TREM2* KO iPSC lines described above, we used the iCell Microglia 01279 line (WT; FUJIFILM Cellular Dynamics Inc. Catalog # R113). The iCell Microglia were thawed and maintained in culture following the manufacturer's guidelines before experiments.

**Flow Cytometry**

iMGL cells were suspended in FACS buffer (DPBS, 2% BSA, 50 μM EGTA) and treated with FC block (1:50, BD Biosciences 564219) for 20 min at 4 °C. Cells were centrifuged and resuspended in FACS buffer before adding CD11b-PE/Cy5 (1:100, Biolegend 101209), CD45-BV650 (1:50, Biolegend 304043), or TREM2-APC (1:50, R&D Systems FAB17291A) and incubated 30 min at 4 °C in the dark. Samples were washed 3× in FACS buffer. Cells were resuspended in FACS buffer and stained with DAPI. Cells were analyzed with the BD Influx Cell Sorter.

**Western Blotting Analysis and Antibodies**

Proteins were resolved via SDS-PAGE using NuPAGE™ 4-12% Bis-Tris gels (Invitrogen) and probed for expression. Proteins were detected using the Odyssey XF Imager (LI-COR Biosciences). Antibodies used included: anti-Phospho-SYK (Cell Signaling), anti-SYK (Cell Signaling), anti-TREM2 (Cell Signaling), and anti-β-actin (Sigma).

**Immunocytochemistry**

iMGL were fixed in 4% paraformaldehyde, permeabilized, and blocked for 1 hr in 10% normal goat serum or 10% fetal bovine serum. Primary antibodies were diluted in 10% normal goat serum containing 0.1% Triton X-100 and incubated overnight at 4°C. Primary antibodies included anti-TREM2 (R&D Systems), anti-IBA1 (Fujifilm), and anti-TMEM119 (Novus)). Following incubation of the primary antibody, cells were washed and incubated for 1 hr at room temperature in Alexa Fluor 568- or 488-conjugated secondary antibody (Invitrogen). Immunostained cells were mounted using VECTASHIELD Mounting Medium (Vector Laboratories) and imaged using the Nikon C1 Digital Confocal System.

**Synaptosome Preparation**

The human temporal lobe was homogenized using a Dounce tissue grinder in homogenization buffer (0.032 M Sucrose, 0.5 mM $CaCl_2$, 1 mM $MgCl_2$, 1 mM $NaHCO_3$ supplemented with protease/phosphatase inhibitor) and centrifuged at 1400×g. The supernatant was saved (S1), and the resulting pellet was re-homogenized in homogenization buffer and centrifuged at 700×g, resulting in the supernatant, S1'. Supernatants S1 and S1' were mixed and centrifuged at 700×g resulting in the supernatant, S2. S2 was centrifuged at 14000×g. The resulting pellet containing synaptosomes was resuspended in buffer (0.32 M Sucrose, 1 mM $NaHCO_3$).

**Oligomer and fibril formation**

Aβ42 oligomers and fibrils were prepared according to the established protocol by Stine *et al. (66)*. Briefly, Aβ42 (Anaspec) was resuspended to 1 mM solution in HFIP. Aliquots were evaporated in a fume hood for 2 hrs, and the resulting peptide film was dried under vacuum in a SpeedVac and stored at -20°C. Immediately prior to use, the films were solubilized to 1 mM in anhydrous DMSO, sonicated in a bath sonicator for 10 min, and diluted to 100 µM in oligomer-forming conditions (cold PBS; incubation at 4°C for 24 h) or fibril-forming conditions (10mM HCl; incubation at 37°C for 24 h).

**pHrodo Labeling**

pHrodo Red (Invitrogen) labeling protocol was adapted from the manufacturer's instructions. Briefly, Aβ42 oligomer or fibril solutions (100 µM) were centrifuged at 16000×g for 2 min to collect the aggregates. Aggregates were washed in 1 mL HBSS and centrifuged for 2 min. The supernatants were aspirated, and 200 µl of pellet aggregates were resuspended by adding 200 µl 0.1M $NaHCO_3$ to the pellets. The pHrodo Red dye (10 mg/ml) was then added per the manufacturer's instructions. pHrodo labeling was performed and incubated for 1 hr in the dark, followed by centrifugation at 16000×g for 2 min. Aggregates were washed 3x with HBSS and resuspended to 100 µM in HBSS.

**Phagocytosis assay**

iMGL were plated at a density of $3x10^5$ cells per well in 96-well plates and incubated for 24 hrs. iMGL were treated with 10 µM SEMA6D in media supplemented with either 30 µg/ml pHrodo Red tagged synaptosomes, 1 mM pHrodo Red tagged Aβ fibril or 400 nM Aβ oligomer and incubated for 24 hrs. Cells were imaged using an IncuCyte live cell imaging system. Analysis was performed by Incucyte Zoom Software. The phagocytic activity of iMGL was quantified using the Incucyte® SX5 Live-Cell Analysis instrument**.** Total cell counts were performed using the CellQuanti-Blue™ Cell Viability Assay kit (BioAssay Systems). Three independent experiments were performed, where all four experimental conditions (WT, KO, WT+SEMA6D, and KO+SEMA6D) were assayed in triplicate (three wells) per experiment. To obtain significance values for Fig. 3B, we used the sumz function from the R metap package (v. 1.8).

**Cytokine assay**

iMGL were plated at a density of $3x10^5$ cells per well in 96-well plates and incubated for 24 hrs. iMGL were treated with 10 µM SEMA6D or 1uM LPS for 24 hrs. Cell media was harvested, and cytokines were quantified using the V-PLEX Human Proinflammatory Panel II (4-Plex) (MSD) according to the manufacturer's instructions. Total cell counts were performed using the CellQuanti-Blue™ Cell Viability Assay kit. Because these experiments were performed in three batches, we analyzed the effects of SEMA6D treatment in cytokine secretion using a linear regression model, including a covariate for the experiment batch.

**iMGL RNA-seq library preparation**

WT or TREM2 KO iMGL were treated with 10 μM SEMA6D for 24 hrs. mRNA was extracted from iMGL using RNeasy Mini kit (Qiagen) following the manufacturer's instructions.

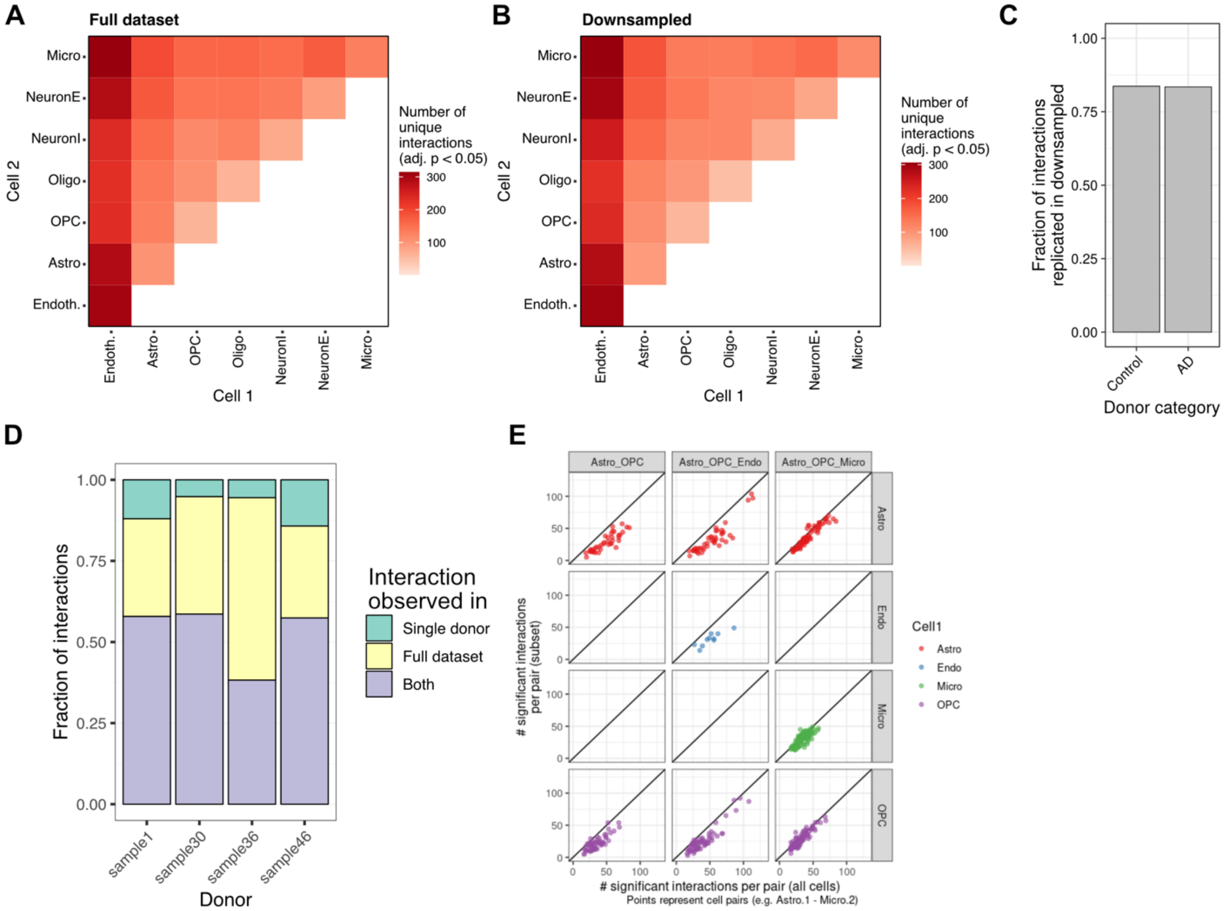


**Fig. S1.** Overview of CellPhoneDB results on the full dataset (**A**) versus a heavily downsampled dataset (maximum 100 nuclei per snRNA-seq cluster) (**B**). **C)** Fraction of replicated CPDB interactions in downsampled dataset compared to the full dataset. **D)** Replication of CPDB interactions detected in the full dataset versus running CPDB in different donors separately. For each donor, only interactions from the corresponding donor category were compared (*e.g.,* sample 1 versus all AD donors). **E)** Number of interactions observed per cell type when comparing the full data (x-axis) versus CPDB using only the cell types indicated in the facet labels (y-axis).

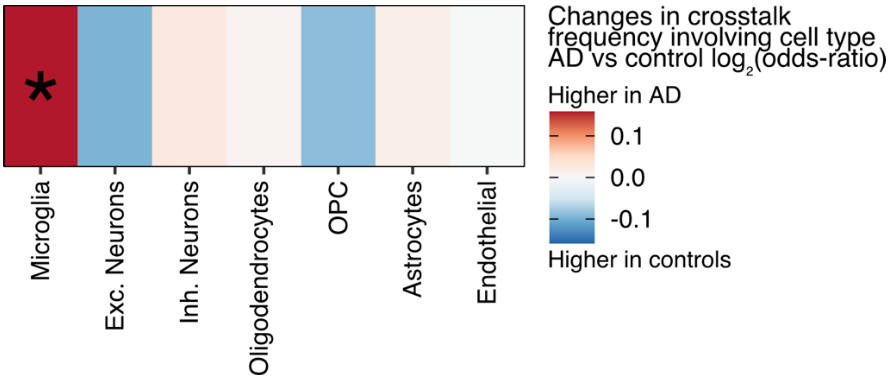


**Fig. S2.** Global changes in crosstalk frequencies per cell type in AD donors versus controls. Related to the heatmap in Fig. 1B.

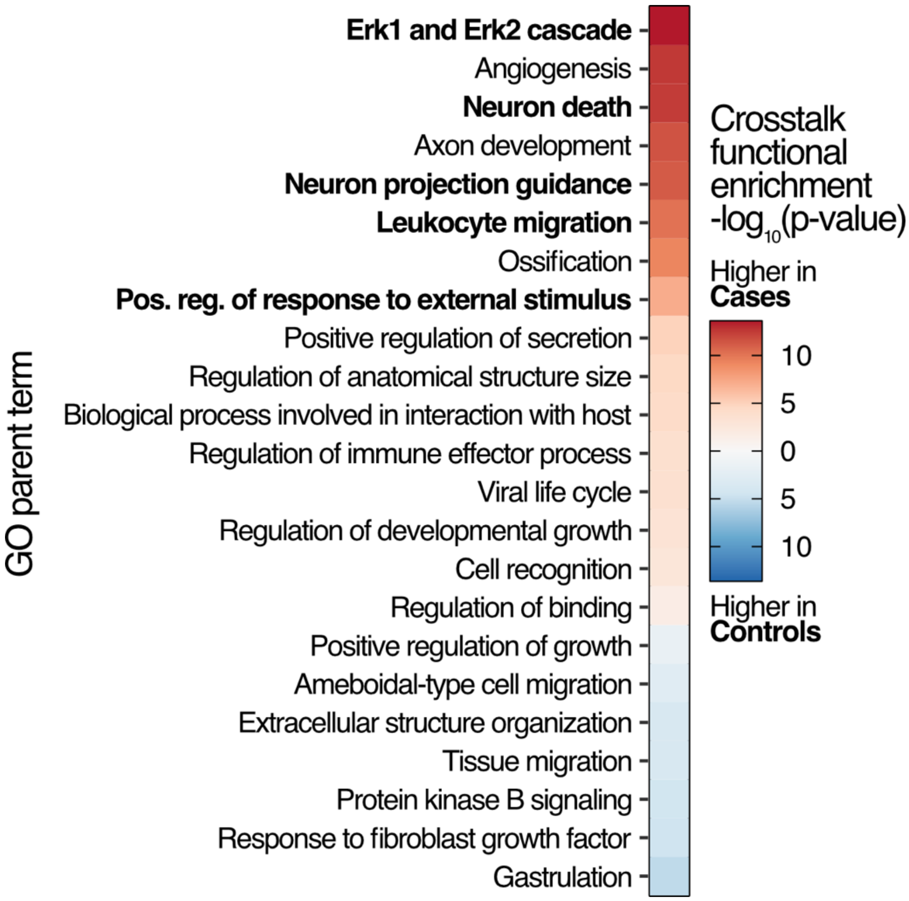


**Fig. S3.** Gene ontology enrichments of the genes associated with crosstalk interactions in cases versus controls.

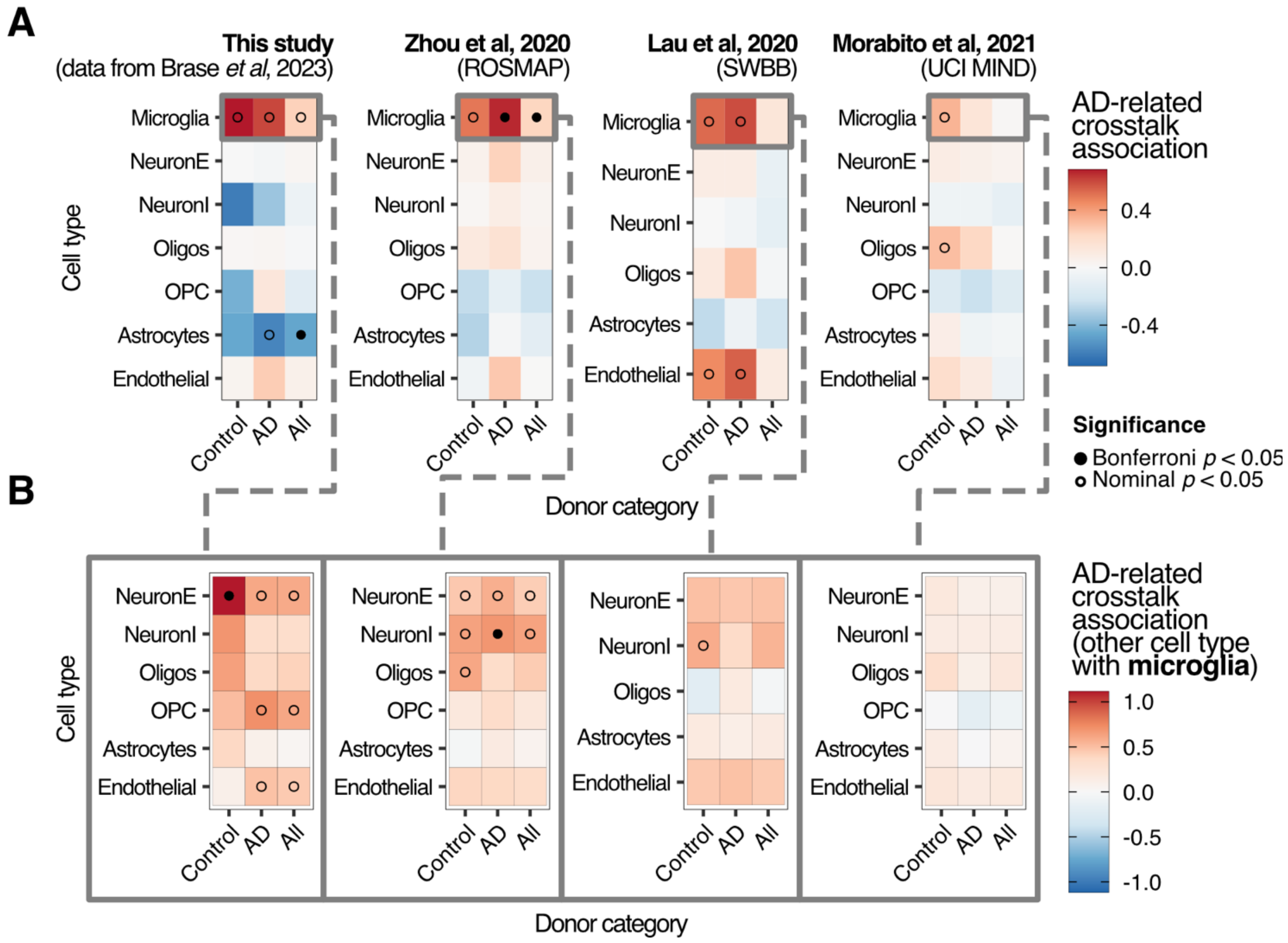


**Fig. S4. A)** Enrichment of AD crosstalk interactions involving each cell type in each donor subset for different snRNA-seq datasets. **B)** Corresponding enrichment of AD crosstalk interactions between microglia and other cell types.

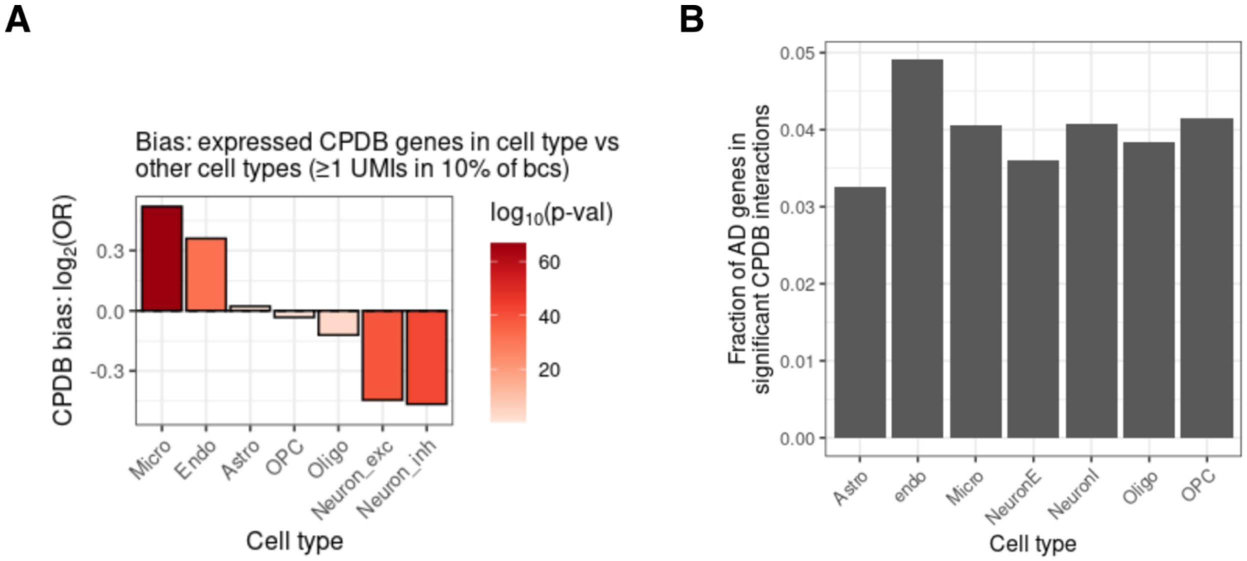


**Fig. S5. A)** Bias of expressed genes in each cell type belonging to a CPDB interaction. This represents how likely we are to encounter interactions involving genes expressed in each cell type in the CPDB database. **B)** Fraction of AD-related genes expressed in each cell type that participate in any CPDB interaction. While we observe a high fraction of genes expressed in microglia involved in CDPB interactions, the representation of AD-related genes is comparable to other cell types (except for endothelial cells).

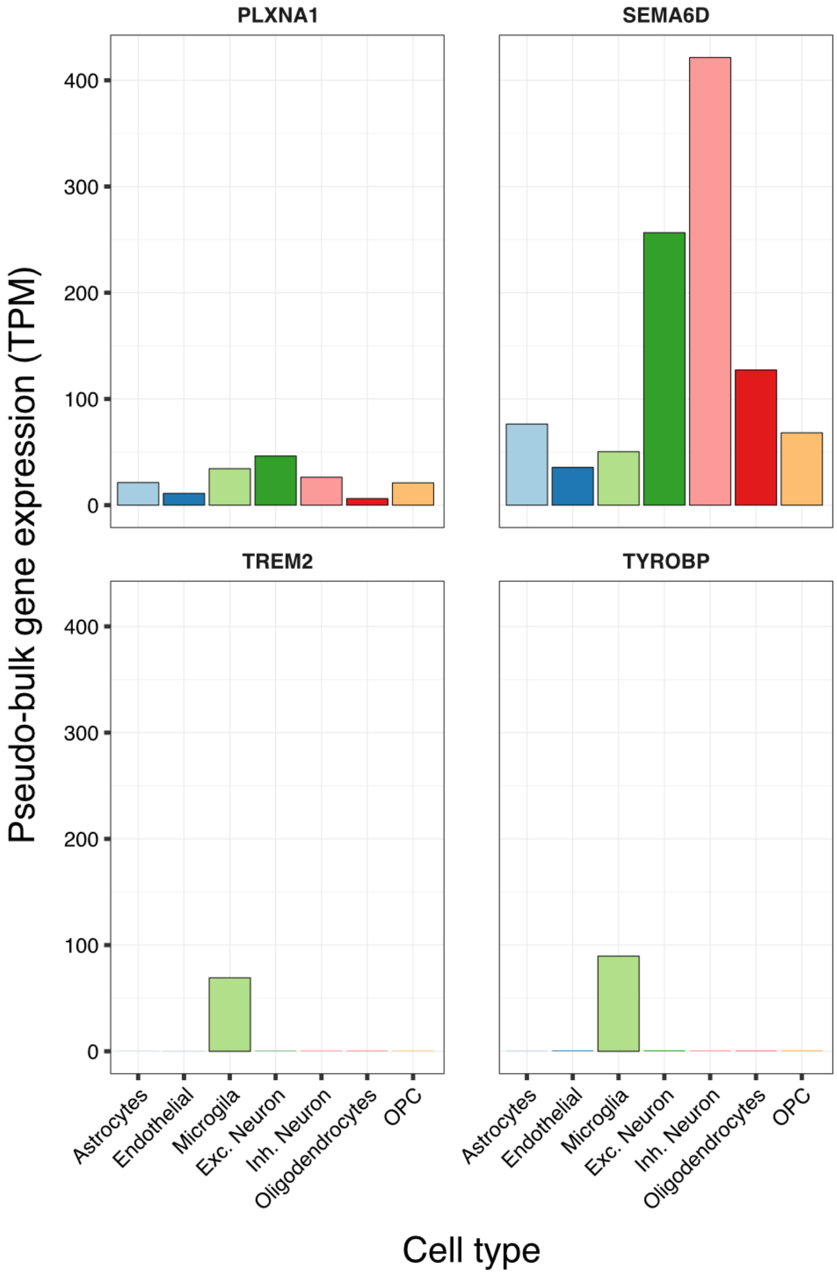


**Fig. S6.** Gene expression patterns of the TREM2-SEMA6D crosstalk interaction components. TPM, transcripts per million.

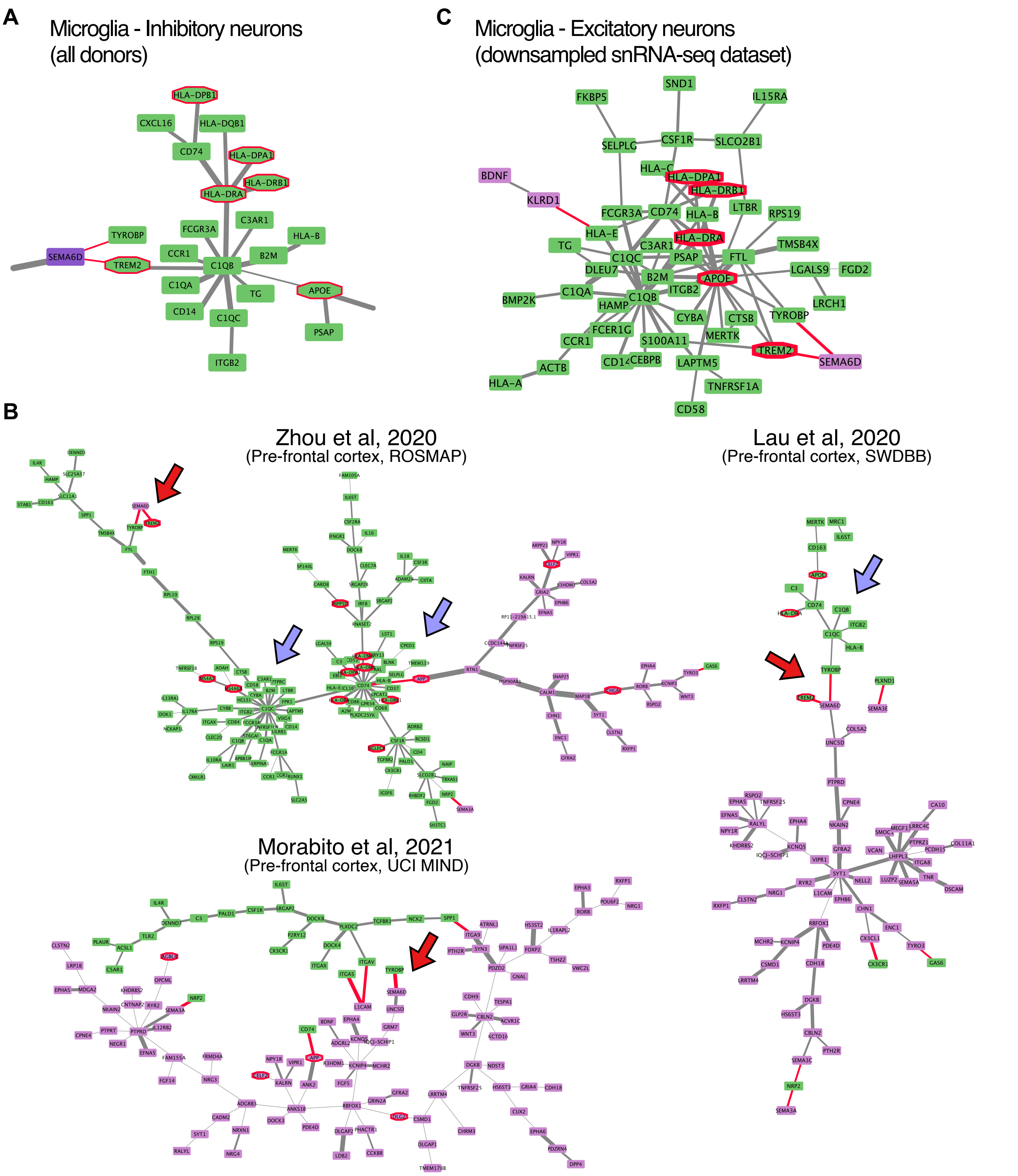


**Fig. S7. A)** Subset of reconstructed inhibitory neurons and microglia crosstalk network using CytoTalk highlighting the *TREM2* sub-network. **B)** Reconstructed excitatory neurons and microglia crosstalk networks using CytoTalk in additional snRNA-seq datasets. Red arrows indicate the TREM2-SEMA6D crosstalk axis, and blue arrows indicate members of the *TREM2* sub-network identified in our data. **C)** Reconstructed *TREM2* crosstalk sub-network with CytoTalk by downsampling the number of nuclei to the donor category with the smallest number of nuclei.

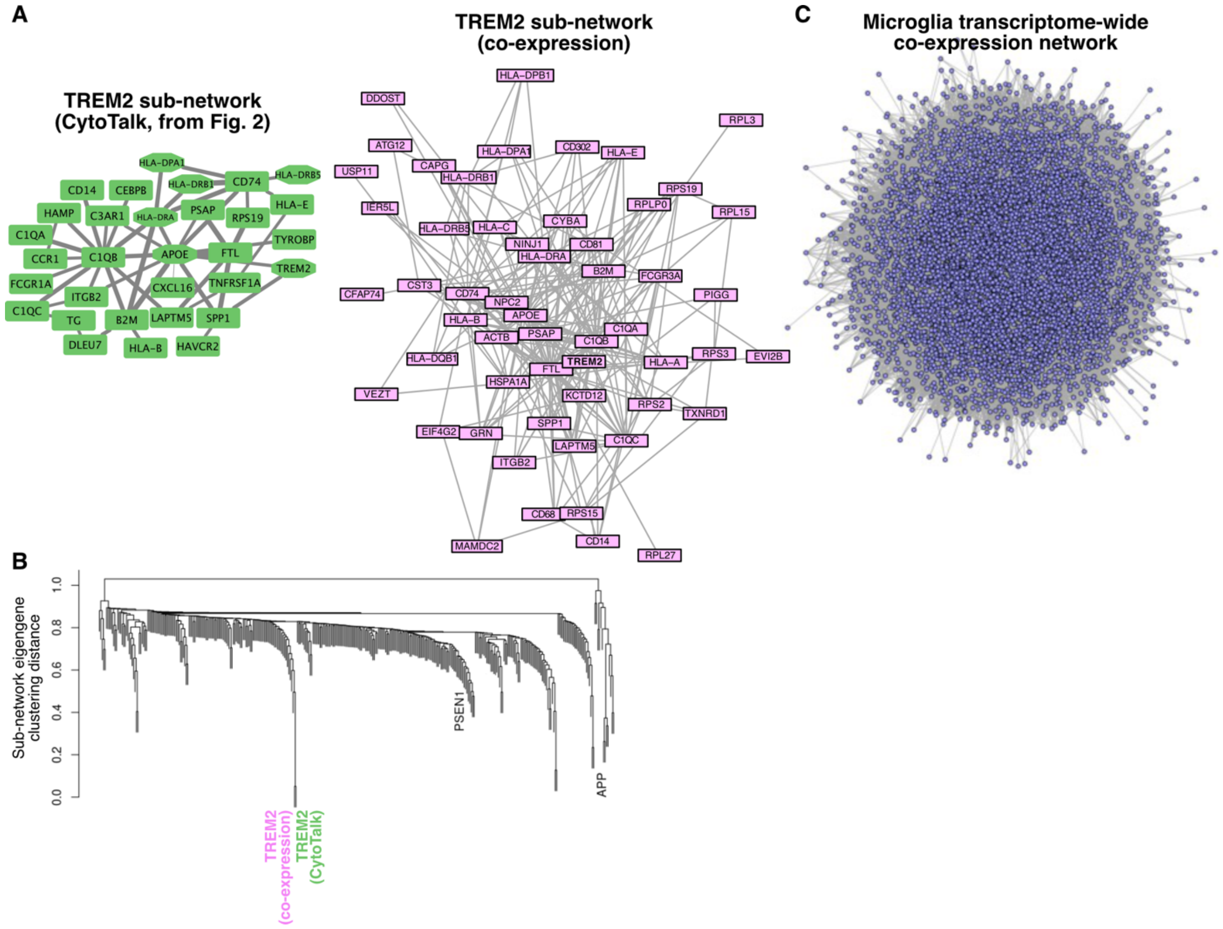


**Fig. S8. A)** Comparison of the TREM2 subnetwork prioritized by CytoTalk and identified directly from the transcriptome-wide microglia co-expression network (C). **B)** Unsupervised clustering of the first principal component of gene expression(eigengene) of all microglia co-expression sub-networks (n = 360; same as Fig. 2H) together with the *TREM2* sub-network identified from CytoTalk. The branch containing the two sub-networks is highlighted on the left of the panel. **C)** Transcriptome-wide (all expressed genes) microglia co-expression network.

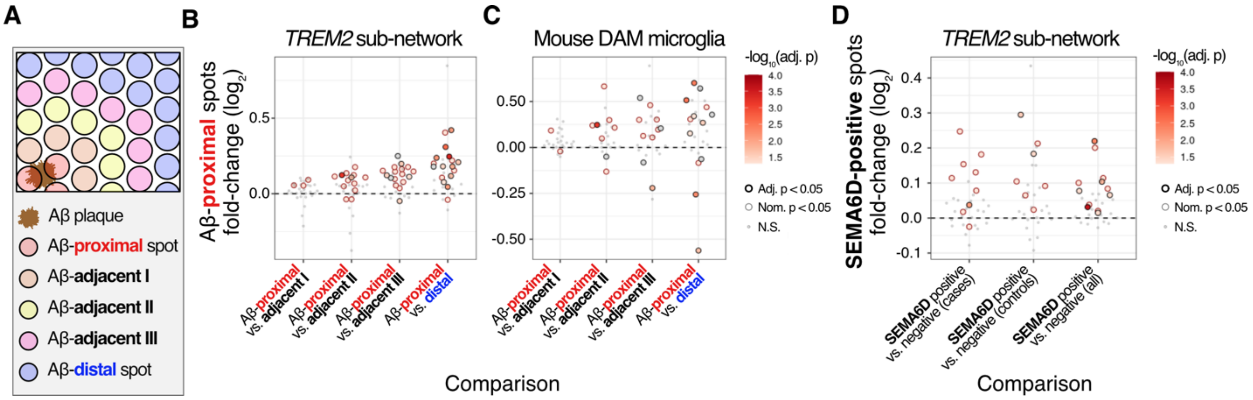


**Fig. S9. A)** Schematic illustrating how Visium spots are classified regarding distance to Aβ plaques for comparisons in (B) and (C). **B)** Transcriptional effects associated with Aβ plaque proximity in the *TREM2* sub-network. Fold changes in (B) and (C) relate to Aβ plaque-proximal spots. **C)** Similar to (B) but comparing signatures of Aβ-associated microglia in mouse single-cell RNA-seq studies. **D)** Comparison of transcriptional effects in the *TREM2* sub-network regarding SEMA6D-positive versus negative spots in different subsets of the data (cases alone, controls alone, and both; the latter corresponds to the bottom row of Fig. 2G).

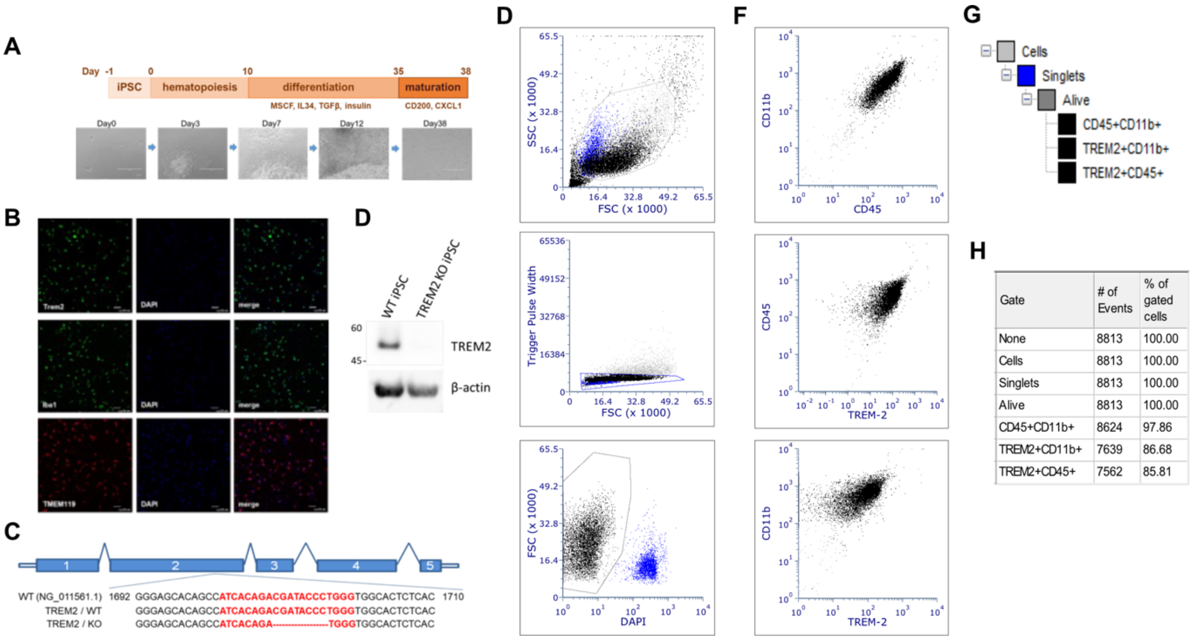


**Fig. S10. A)** Schematic of the differentiation procedure and brightfield imaging of iPSC-derived microglia (iMGL). **B)** mature iMGL stained with microglial markers, TREM2, IBA1, and TMEM119 and co-stained with DAPI. **C)** Schematic of DNA (red) targeted by gRNA against TREM2 for CRISPR/Cas9 modification and showing an 8-bp homozygous deletion within exon 2 of the TREM2 gene. **D)** Western blot analysis of wild-type (WT) and TREM2 KO iPS cell lysates immunostained for TREM2 with β-actin as the loading control. **E)** Flow cytometry gating strategy based on SSC, FSC (upper panel). Doublets excluded based on pulse width, FSC (middle panel). Live cells were identified based on FSC and DAPI (bottom panel). **F)** Identification of gated microglia based on the expression of CD11b, CD45 (upper panel); CD45, TREM2 (middle panel); CD11b, TREM2 (bottom panel). **G)** Diagram of gating strategy from (E) and (F). **H)** Gating strategy: number of events and percent of gated cells from (E) and (F).

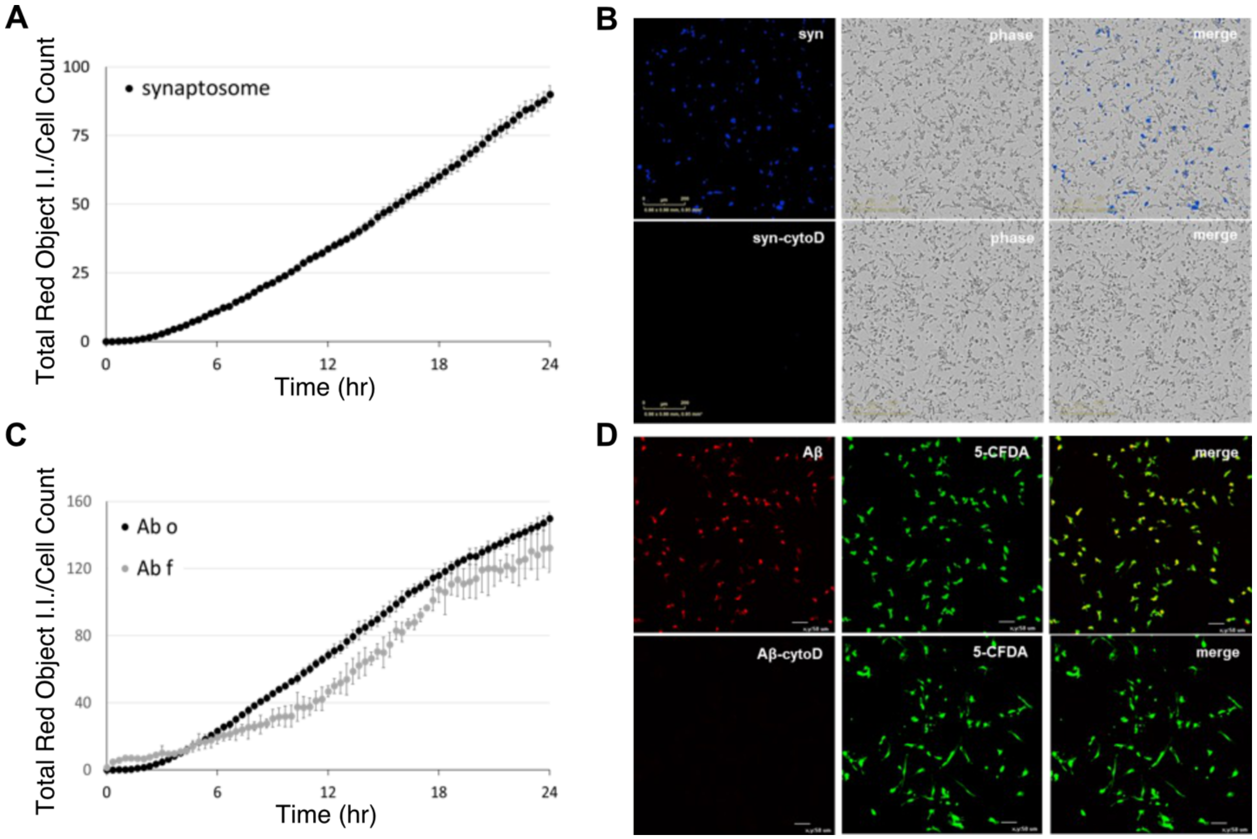


**Fig. S11. A)** Phagocytosis quantification using Incucyte live-cell analysis system on iMGL treated with pHrodo-labeled synaptosomes (syn). **B)** Fluorescent and brightfield microscopy of iMGL treated with pHrodo-labeled synaptosomes (syn) with/out cytochalasin D. **C)** Phagocytosis quantification using Incucyte live-cell analysis system on iMGL treated with pHrodo-labeled Aβ oligomer (Abo) and Aβ fibril (Abf). **D)** Fluorescent microscopy of iMGL treated with pHrodo-labeled Aβo with/out cytochalasin D and co-stained with 5-CFDA membrane marker. I.I., integrated intensity.

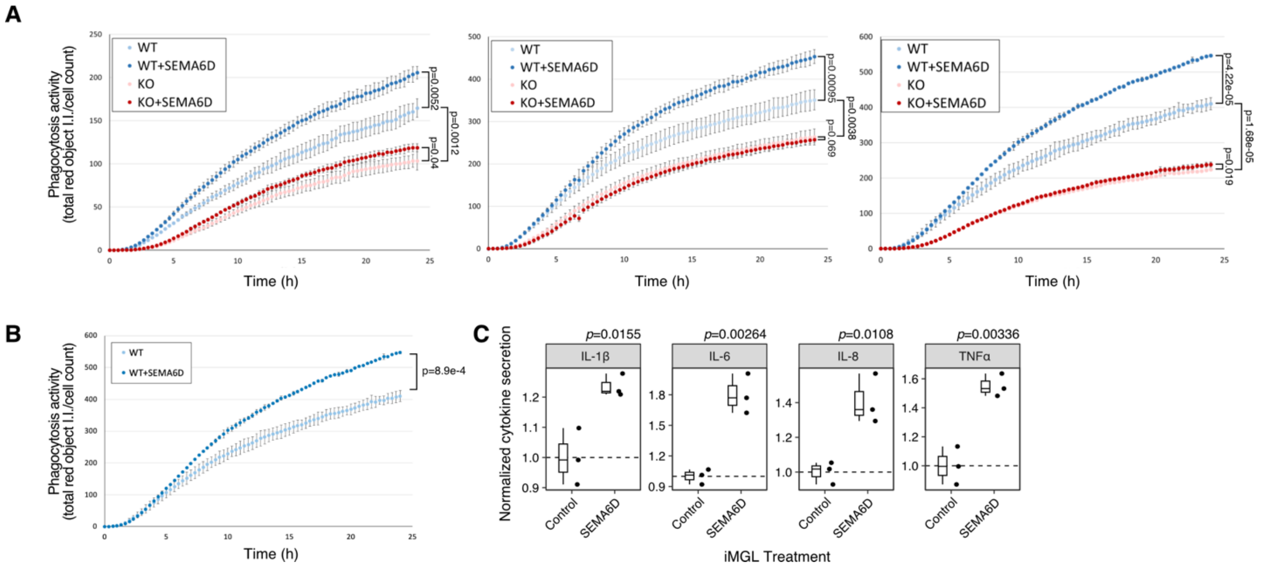


**Fig. S12. A)** Phagocytosis of synaptosomes by WT or *TREM2* KO iMGL treated with SEMA6D (5 µg/ml), related to Figs 3a-b. Three independent experiments; mean ± SD values. Per experiment *p*-values were calculated using two-sample t-tests. I.I., integrated intensity. **B)** Similar to (A) for an additional WT iMGL line. **C)** Quantification of media from an additional WT iMGL line treated with SEMA6D (10 µg/ml) for cytokine release using the Mesoscale V-plex neuroinflammation panel. Boxplots indicate median and interquartile ranges, p-values calculated using linear regression.

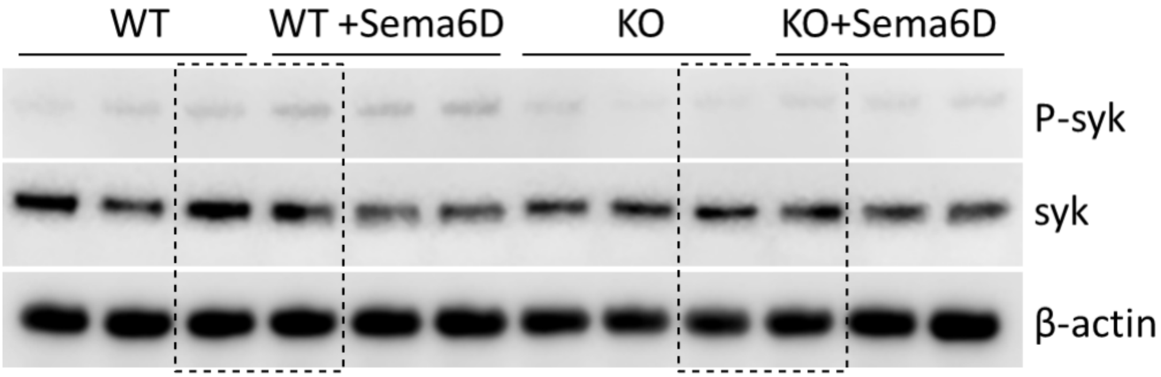


**Fig. S13.** Individual replicates for the western blot analysis of p-SYK and total SYK in WT and *TREM2* KO iMGL treated with SEMA6D (10 µg/ml). β-actin as the loading control. Related to Figs. 3d-e. Dashed boxes indicate sections used for Fig. 3D.

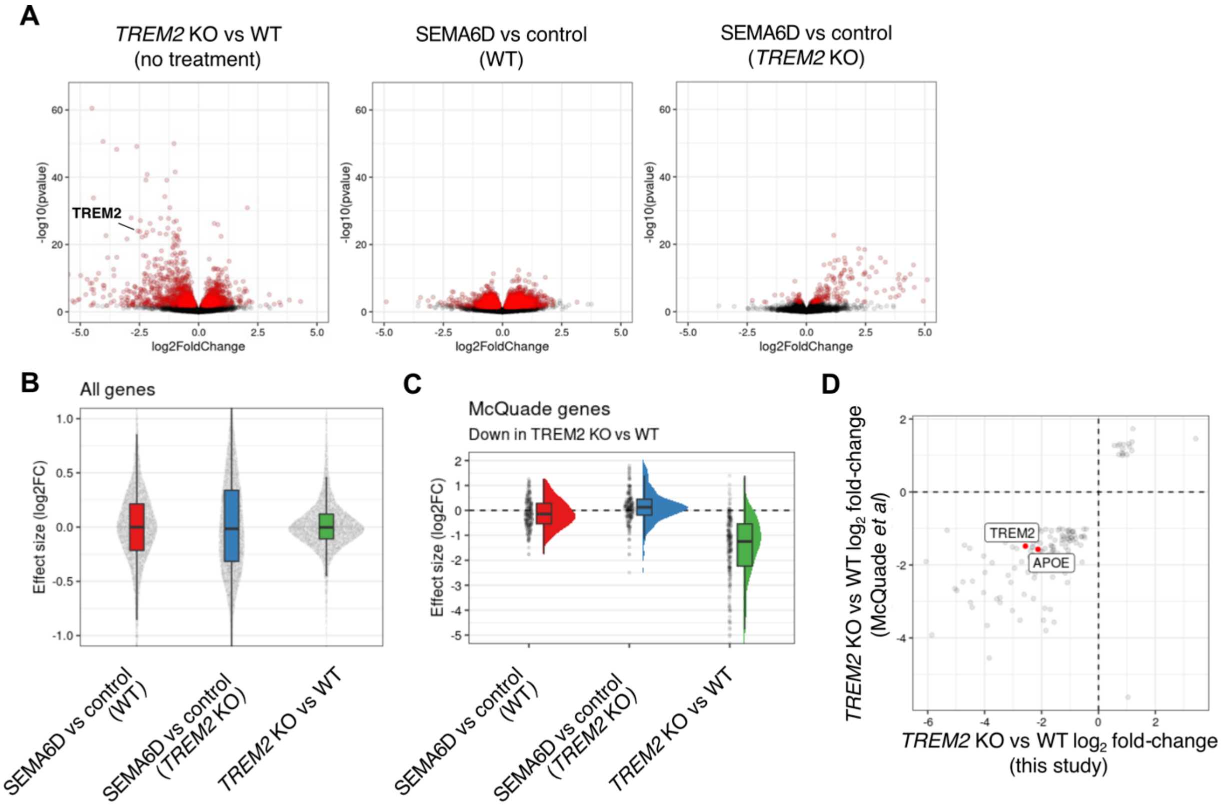


**Fig. S14. A)** Volcano plots for the differentially expressed genes in different iMGL RNA-seq experiments. Red genes correspond to differentially expressed at 10% FDR. **B)** Effect size distribution of the three iMGL experiments. **C)** Effect size distribution of downregulated genes in TREM2 KO iMGL from the McQuade et al. 2018 study in the three iMGL RNA-seq experiments. **D)** Comparison of reported effect sizes from the McQuade study and the TREM2 KO vs. WT iMGL RNA-seq experiment from this study.
